# SpatialCCCbench: Standardized Metrics for the Systematic Evaluation of Spatial Cell-Cell Communication Methods

**DOI:** 10.64898/2026.05.19.724475

**Authors:** Qian Wang, Dawei Liu, Yuan-Yuan Li, Wentao Dai

## Abstract

Spatial transcriptomics (ST) enables transcriptome profiling with preserved spatial context, providing spatial dimensions that are essential for understanding complex intercellular signals in tissue architecture. ST-based CCC tools integrate spatial and molecular information to decipher intercellular interactions from a spatially informed perspective. Despite the rapid evolution of many CCC computational tools, a systematic assessment of their performance in handling ST-specific heterogeneity, utilizing spatial information efficiently, and robustness against technical or biological noise is still lacking. To address this gap, SpatialCCCbench incorporates classification accuracy, spatial signal features, robustness, and user-friendliness, aiming to guide the selection of optimal CCC inference tools across diverse spatial biology contexts. SpatialCCCbench systematically evaluates the scenario-specific applicability of ST-based CCC tools. It helps users select tools according to their analytical objectives and provides a practical benchmark for future method development.

**Highlights:**

1. Established a multi-dimensional benchmark suite to evaluate cell-cell communication (CCC) inference methods in spatial transcriptomics.
2. Characterized the spatial patterns of CCC signals across diverse tissues using spatial autocorrelation and local diversity analysis.
3. Systematically assessed the robustness of CCC inference tools across six common experimental noise scenarios in spatial transcriptomics.
4. Integrated boundary-feature analysis, a mechanistically important component for biological interpretation, to uncover spatial preferences and algorithmic biases in CCC methods.
5. Provided guidelines to assist in the selection of optimal CCC inference tools tailored to various spatial biology contexts.

**Graphic Abstract:** 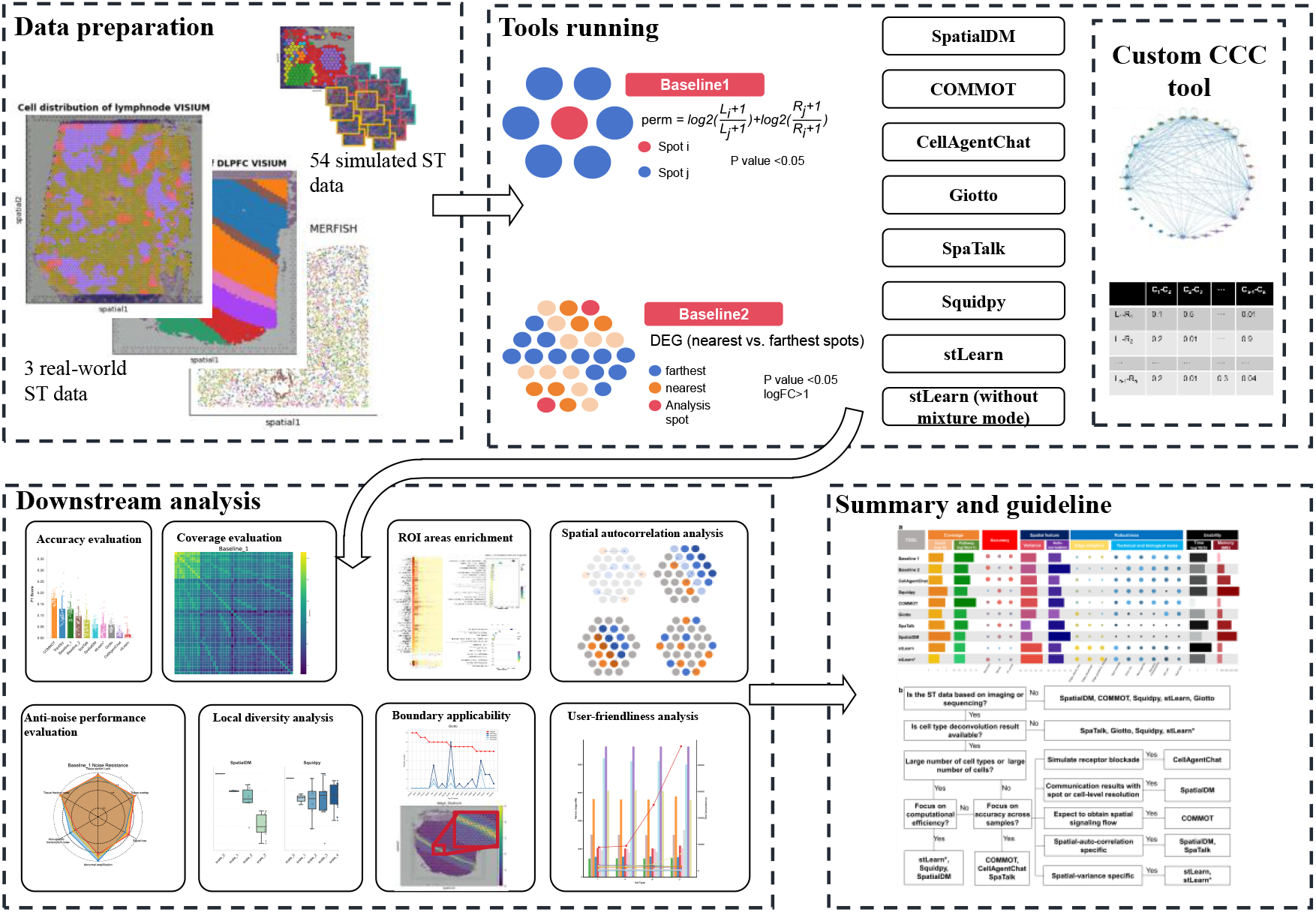

## Introduction

The complex biological activities of multicellular organisms are fundamentally governed by complex tissue architectures and diverse cell subpopulations that function within specific niches[1, 2]. Limitations in analyzing cellular heterogeneity and communication at the single-cell level have been significantly addressed by the development of spatial transcriptomics (ST), which enables transcriptome-wide profiling while preserving native tissue coordinates[3-5]. Furthermore, the spatial accuracy and sensitivity of these profiles have facilitated the identification of transcriptomically distinct subpopulations and ecotypes in tumor microenvironments and specific pathological characteristics[6-10]. Consequently, the rapid evolution of ST has opened up a transformative analytical perspective for mining complex organizational information and characterizing tissue-level features[11].

ST has provided a powerful approach for deciphering intercellular signaling and mapping interaction hot spots within tissues, offering unprecedented spatial context for understanding cell-cell communication (CCC)[12, 13]. As a fundamental process, CCC orchestrates development, maintains tissue homeostasis, and regulates physiological function in multicellular organisms, while its dysregulation is also central to the mechanisms of many complex diseases[14, 15]. In conventional single-cell studies, although the co-expression of ligand-receptor (LR) pairs can be inferred, the absence of physical spatial information often restricts such analyses to non-spatial contexts. The spatial proximity of ligands and receptors from ST provides more robust evidence for biological interactions compared to traditional single-cell RNA-seq (scRNA-seq) methods[16, 17]. Consequently, a multitude of computational tools have been developed to model these localized signaling events, employing diverse mathematical frameworks ranging from optimal transport and spatial graphs to deep learning-based foundation models[18-21]. However, the performance of these algorithms is significantly influenced by technical factors such as molecular diffusion, spatial information preference, and noise interference[22, 23].

In recent years, a number of researchers have systematically integrated and evaluated the performance of various CCC tools on extensive datasets, providing informative metrics and reliable guidance for the selection of CCC analysis tools. For instance, Liu et al. benchmarked 16 interaction methods by integrating scRNA-seq with ST data and analyzed CCC signals by categorizing interactions into short- and long-range effects based on spatial distance distributions in ST data[24]. Simultaneously, the ESICCC framework evaluated prior interaction information, LR scoring algorithms, and intracellular signaling complexity to provide a selection guide for scRNA-seq-based and ST-based tools, while peripherally assessing the impact of incorporating spatial data[25]. Despite the prevalence of scRNA-seq-centered evaluation metrics, a systematic assessment of how tools capture ST-specific heterogeneity and utilize spatial coordinates remains critically lacking. Furthermore, emerging ST-based LR inference methods, such as Giotto[26], stLearn[15], COMMOT[18], and SpatialDM[16], still require independent benchmarking strategies that account for their specific spatial logic, which cannot be comprehensively evaluated by existing frameworks.

Therefore, a systematic comparison of existing CCC inference tools is urgently required to establish standardized evaluation metrics and provide guidance for the selection of optimal analytical strategies in spatial biology. Here, we present SpatialCCCbench, a comprehensive benchmarking framework that integrates multi-dimensional evaluation criteria, to guide the selection of optimal CCC tools across diverse spatial contexts.

## Results

A systematic and comprehensive benchmarking framework is presented to evaluate the signal features derived from tools for CCC analysis in ST data. Three real-world ST datasets, six noise-simulated ST datasets, three boundary-simulated ST datasets, and 45 simulated heterogeneous datasets were used to evaluate 10 methods with two baselines (see Methods and Fig. S1). This framework was designed to characterize CCC signals from multiple and individual ST samples, with systematic analysis encompassing classification accuracy, signal coverage, biological relevance, robustness, spatial autocorrelation, local signal diversity, reliability of boundary features and user-friendliness, as demonstrated below.

### Classification accuracy of CCC signals on multiple or individual samples

Focusing on tissue heterogeneity across pathological samples, we evaluated CCC classification accuracy on simulated ST datasets and real-world ST datasets using precision, recall, and F1 score as described in the Methods. As shown in Additional file 1: Fig. S2a, b, COMMOT achieved the best overall balance, with higher precision and F1 score than both Baselines, while Squidpy and CellAgentChat showed higher recall. stLearn varied more across samples, suggesting sensitivity to tissue-specific spatial patterns.

Because CCC signals aggregated from multiple samples may be biologically inconsistent, we further assessed tool performance across platforms and tissues. COMMOT and stLearn performed best on Visium lymph node data (Fig. S2c, d, e), SpatialDM and the baselines were stronger on multiplexed error-robust fluorescence in situ hybridization (MERFISH) mouse brain data (Fig. S2f, g, h). In addition, Squidpy and stLearn ranked higher on dorsolateral prefrontal cortex (DLPFC) slices (Fig. S2i, j, k). These results indicate that CCC tool performance varies substantially across tissues and platforms, and appropriate tool selection is essential for reliable biological interpretation.

### Coverage and similarity of LR pairs maintaining CCC across tools

ST techniques suffer from low detection rates for low-expression genes, impacting CCC inference. Using an empirical LR-cell-cell pattern to quantify signal coverage, we observed that baselines constructed solely through simple LR co-localization and cell-type mapping capture only a subset of interactions. Compared to the output of more sophisticated CCC tools, the interaction yield of these baselines remains relatively limited. Specifically, we found that SpatialDM and stLearn captured the most interactions from specified cell types, while CellAgentChat used stricter filtering with nearly consistent interaction counts across samples (see Fig. S3a, b). In addition, the enrichment signals generated by these genes also showed distinct features, with Squidpy achieving broader coverage in the enrichment of characteristic signaling (as shown in Fig. S4). Across these large-scale interaction patterns, tools generally showed higher Similarity indices[25] and lower Jaccard distances (Fig. S3d). However, substantial heterogeneity remained, and the low overlap between tools suggests distinct capture preferences and spatial variability. Simple similarity metrics are thus insufficient to capture complex CCC networks, necessitating a more robust evaluation framework to address these tool-specific biases and spatial complexities.

### Spatial autocorrelation analysis of CCC tools

ST inherently captures the spatial organization of tissues, where genes exhibit distinct spatial clustering and co-expression characteristics driven by cell-type specificity and biological regulatory mechanisms. Because these spatial features directly influence CCC signal detection, evaluating CCC tools requires a robust classification of spatial signaling preferences. To achieve this, we established a comprehensive workflow integrating a Gini coefficient, Moran’s I, and Geary’s C to describe spatial autocorrelation. To rigorously evaluate interactions, Exploratory Spatial Data Analysis (ESDA) was employed to assess the spatial autocorrelation of individual genes[27, 28]. For LR spatial correlation, we evaluated the integrated expression patterns of LR pairs using modified Moran’s I and Geary’s C statistics, as detailed in the Methods and Fig. S5. As shown in Fig. 1a and b, local Moran’s I measures the spatial clustering characteristics of gene distribution, whereas local Geary’s C assesses variance across local spatial spots.

**Fig. 1.**
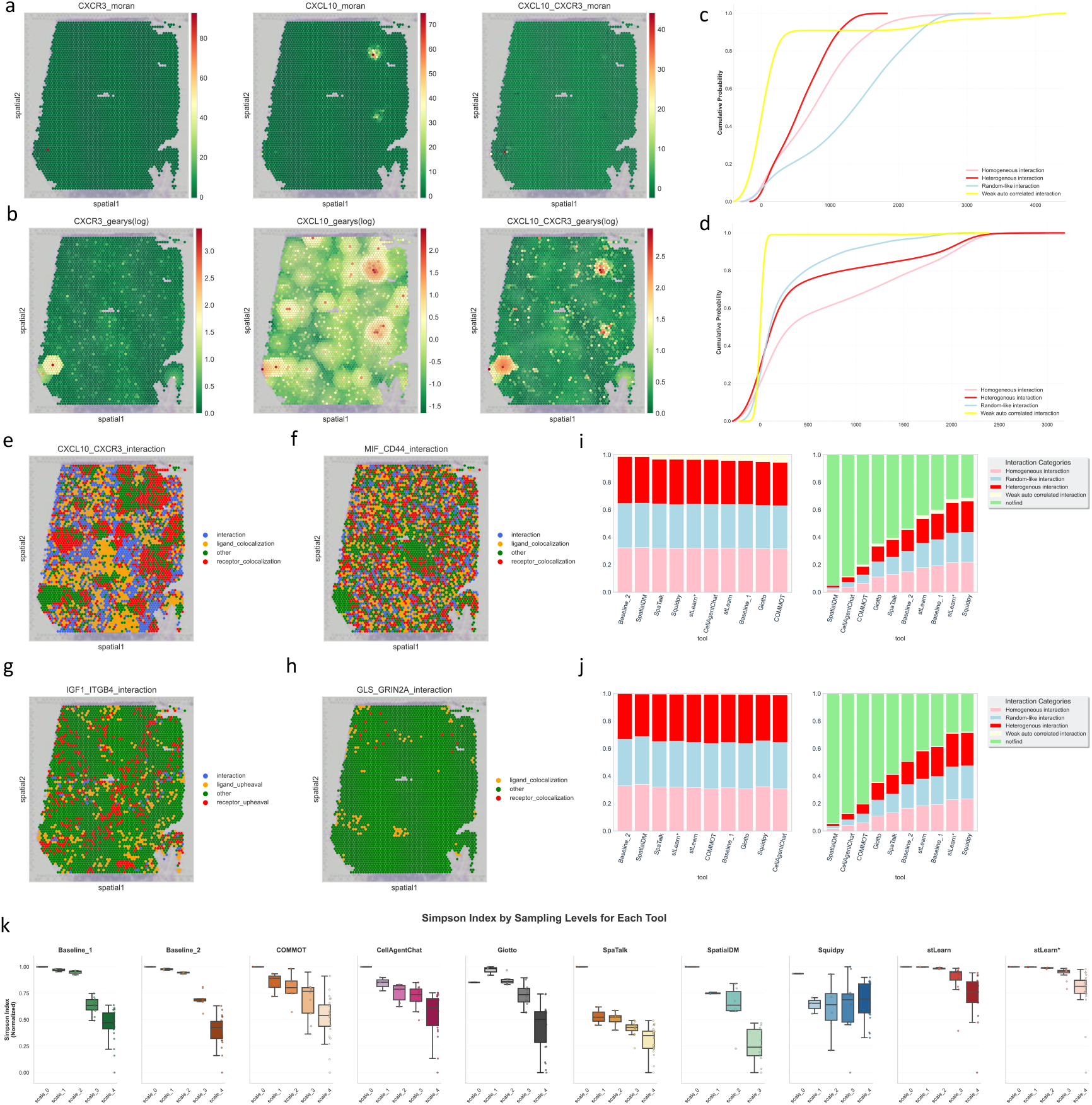
**a–b**. Spatial distribution of local Moran’s I and Geary’s C for interacting single genes and integrated LR pair (CXCL10-CXCR3). **c–d**. Accumulation curves of interaction spot numbers for the four interaction categories defined by autocorrelation analysis using single genes (c) or integrated LR pairs (d). **e–h**. Schematic illustration of homogeneous interaction (e; colocalization with local Moran’s I > 0 and local Geary’s C ∈ (0, 0.5]), random-like interaction (f; colocalization with local Moran’s I > 0 and local Geary’s C ∈ (0.5, 1.5)), heterogeneous interaction (g; local signal transition with local Moran’s I > 0 and local Geary’s C ≥ 1.5), and weakly autocorrelated interaction (h; signal-aggregated colocalization with local Moran’s I > 0 and local Geary’s C ∈ (0, 0.5]) based on single-gene autocorrelation analysis. The weakly spatial colocalization pattern in panel h reflects interaction regions where ligand-receptor signals are locally aggregated and colocalized, rather than regions with weak autocorrelated interaction. **i–j**. Spatial autocorrelation and integrated LR correlation analyses for heterogeneous interaction and contact CCC signals across different tools. **k**. Normalized Simpson index across sampling scales for each CCC tool.

Based on these frameworks, spatial autocorrelation analysis identified four distinct spatial distribution patterns for LR pairs. Homogeneous interaction (Fig. 1e), exemplified by CXCL10-CXCR3, reveals strong regional clustering indicative of specific biological processes such as localized immune chemotaxis. Random-like interaction (Fig. 1f), as seen with MIF-CD44, spans multiple cell types with extensive co-localization distributed broadly throughout most of the tissue regions. A heterogeneous interaction pattern (Fig. 1g), characterized by IGF1-ITGB4, features spatially variable expression zones that typically reflect secretion-mediated intercellular interactions. Weak spatially autocorrelated interactions like GLS-GRIN2A (Fig. 1h) display insufficient spot-level clustering and lack a broader and biologically meaningful spatial macro-organization. These metrics generated cumulative probability curves for interaction regions in Fig.1c and d. The single-gene autocorrelation analysis shown in Fig.1c proved highly sensitive in detecting small-scale heterogeneous interaction and homogeneous interaction patterns, whereas the integrated LR-spatial correlation analysis presented in Fig. 1d, which integrates receptor and ligand expression within the same parameters, is superior at uncovering intuitive random-like interaction features resulting from global LR co-localization.

Finally, we classified the CCC signals identified by various tools using single-gene (Fig. 1i) and integrated LR (Fig. 1j) autocorrelation analysis frameworks to evaluate their spatial informativeness. Baseline 1 can capture specific local signals through small-scale permutation tests, yet its neglect of global features leads to the inclusion of certain weak spatial autocorrelation signals. Conversely, Baseline 2 excludes LR pairs characterized by weak spatial differential distributions and an absence of local clustering because of its stringent filtering of cells in distal regions. This approach establishes a rigorous and complementary filtering principle for spatial autocorrelation within this baseline. SpatialDM demonstrated the most robust performance, with its identified interactions highly concentrated in the biologically informative homogeneous and random-like interaction categories owing to its dual Moran’s I-based calculation module. In contrast, tools such as stLearn, Squidpy, and Baseline 1 detected large numbers of interactions reflected by low “not found” rates, but they exhibited a higher proportion of weakly auto-correlated signals across other methods, indicating that these tools utilize relaxed spatial filtering criteria, capturing many interactions lacking verifiable spatial relevance.

### Spatial variance analysis on local diversity of CCC signals

With the advancement of high-resolution ST technology, the local structure of tissues has been further resolved through increasingly complex, large-scale data. Whether a CCC tool can identify the signal content with local diversity has become a key performance criterion for evaluating tool performance. CCC signals can exhibit considerable diversity and dynamic trends at the sample scale, particularly as tissue heterogeneity and the number of cell types vary as shown in Fig. S3c. The Simpson index was applied to assess the local diversity of LR interactions, where each LR-cell-cell pair was treated as a distinct population (see Methods for details)[29].

Overall, a significant decline in diversity was observed from the whole-tissue level to local subsections, as illustrated in Fig. S3c, e. The Simpson index calculated across sampling scales revealed mild fluctuations among the results from different tools, but a consistent decreasing trend in the normalized Simpson index was evident. As shown in Fig. 1k, Baseline1 and Baseline2 exhibited a pronounced scale-dependent trend attributable to their scan-search mode. Similarly, interactions derived from stLearn, COMMOT, and CellAgentChat varied considerably because of the incorporation of variable spatial elements. These findings indicate that these tools are sensitive to local diversity, supporting the scale effect of the Modifiable Areal Unit Problem (MAUP)[3]. Furthermore, while stLearn sustains a descending gradient, it simultaneously maintains robust local diversity characteristics. Notably, the local diversity estimated by Giotto and Squidpy across the whole tissue (scale 0) was lower than that in the smaller sampling areas, exhibiting an inconsistent downward trend. This pattern potentially reflects a more conservative estimation strategy when these methods are applied to broader tissue regions.

Additionally, high variation in the Simpson index was also detected among the smallest tissue subunits. This observation aligns with the zoning effect of MAUP, which highlights that statistical indices are influenced by the combined effects of gene expression patterns and cell-type composition at a given spatial scale, leading to pronounced diversity differences even within the same sampling stratum. Specifically, Squidpy yielded a higher Simpson index at scale 4 compared to scale 1, suggesting that local regions may retain sufficiently detailed gene interaction information.

### Spatial boundary adaptation of CCC tools

Spatial self-organization, inherent in structures like lymph node germinal centers and tumor niches, establishes distinct boundaries with unique molecular and signaling signatures. Identifying these edge-specific patterns is a central challenge in ST analysis, where performance often diverges between tools emphasizing integrated gene expression and those prioritizing spatial proximity. We systematically assessed CCC tool adaptation at these boundaries using three simulated edge features representing different perturbations. As demonstrated in Fig. S9, this framework evaluates how various computational approaches characterize transitional signaling signatures across distinct tissue domains.

Specifically, as depicted in Fig. 2a, b, Baseline 1, based on neighbor permutation, can be strongly influenced by gradient change and equalization, indicating that permutation on local spots relies heavily on the edge gradient and accurate expression. Additionally, Baseline 2, based on differentially expressed gene (DEG) analysis, is only slightly influenced by edge equalization while maintaining CCC stability under gradient change, suggesting that DEG analysis can preserve the specificity of spatial gene expression. These baselines establish a spectrum of signal detection preferences among tools. For instance, Squidpy and CellAgentChat yielded Pearson correlation coefficients close to 1 while Giotto and COMMOT remain less affected by boundary noise, suggesting robust performance under noisy or variable edge conditions. In contrast, SpaTalk, stLearn, and SpatialDM displayed greater output variance, indicating higher sensitivity to fine-grained morphological changes at tissue edges.

**Fig. 2.**
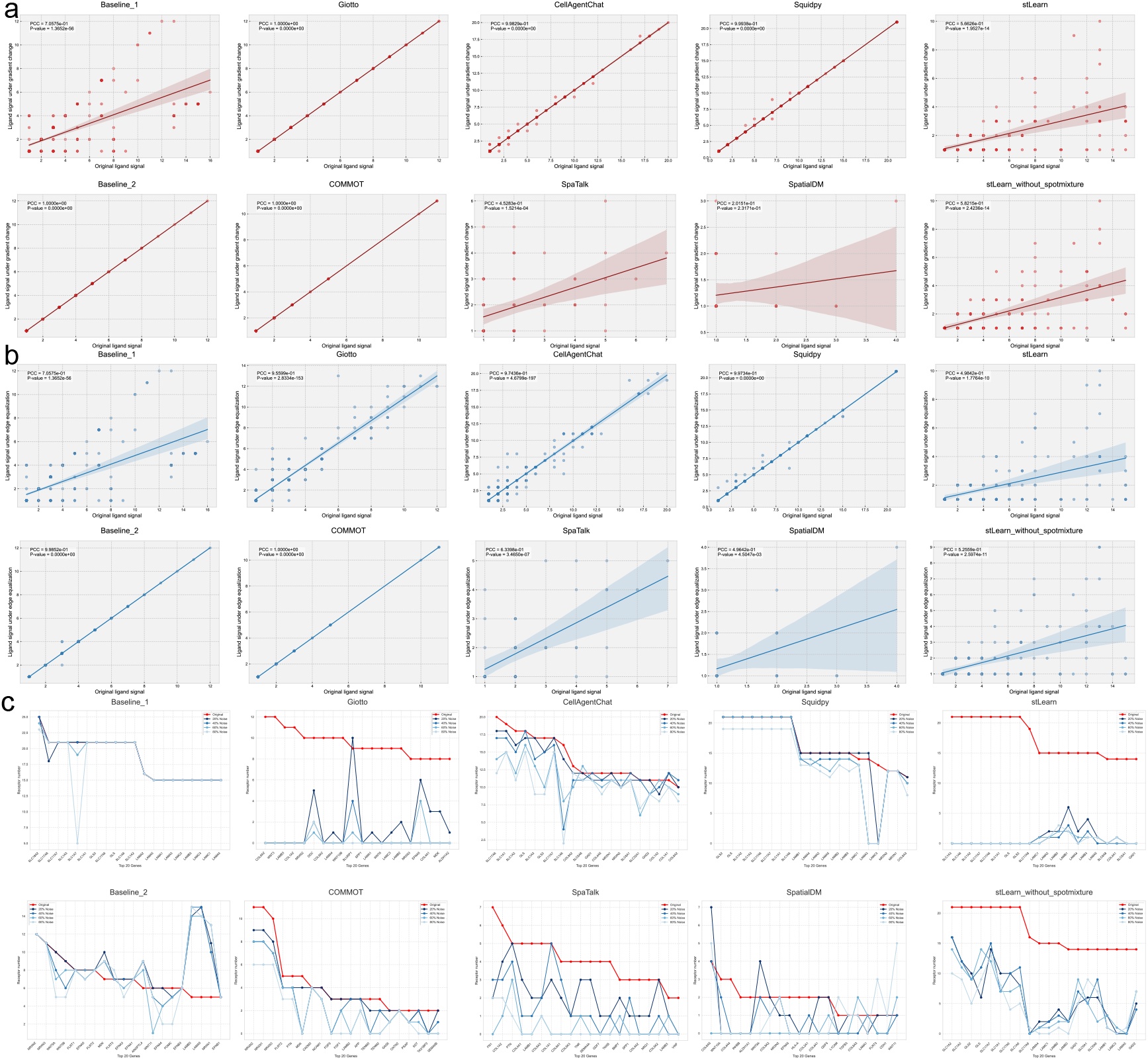
Spatial boundary adaptation of CCC tools under gradient change, edge equalization, and signal loss. **a**. Correlation between CCC signals from the original data and edge-equalized data. **b**. Correlation between CCC signals from the original data and gradient changed data. **c**. Changes in signal fluctuation across different signal-loss levels.

To further examine the impact of gene expression dropout, signal strength changes across four noise levels were recorded, as shown in Fig. 2c. In general, most tools exhibited interaction signal loss, with ligand strength lower than the original level, whereas some tools showed higher ligand signal strength, which indicates noise-induced signal inflation or potential spurious interactions. Specifically, Baseline 2, stLearn, and SpatialDM were easily influenced by signal loss at boundaries, suggesting that their CCC results depend strongly on accurate edge gene expression obtained from high-precision and sensitive ST techniques. In contrast, Squidpy, CellAgentChat, and COMMOT showed strong noise resistance, indicating that these methods focus more on cell-type cluster features than on edge gene expression at the boundary.

### Robustness analysis under technical and biological noise on spatial data

During the sample preparation of spatial transcriptomics data, technical and biological noise is inevitably introduced, primarily due to tissue fragility and limited signal capture from thin sections. Whether CCC inference results are affected by such noise remains a critical consideration in tool selection. To address this, six common noise scenarios were simulated, including tissue section lack, tissue overlap and tissue fracture noise (arising from technical errors in sample preparation), signal loss and abnormal amplification (resulting from transcript capture and sequencing errors), and non-specific transcription noise caused by background biological signals. As shown in Fig. 3a, all tools were evaluated under these six conditions.

**Fig. 3.**
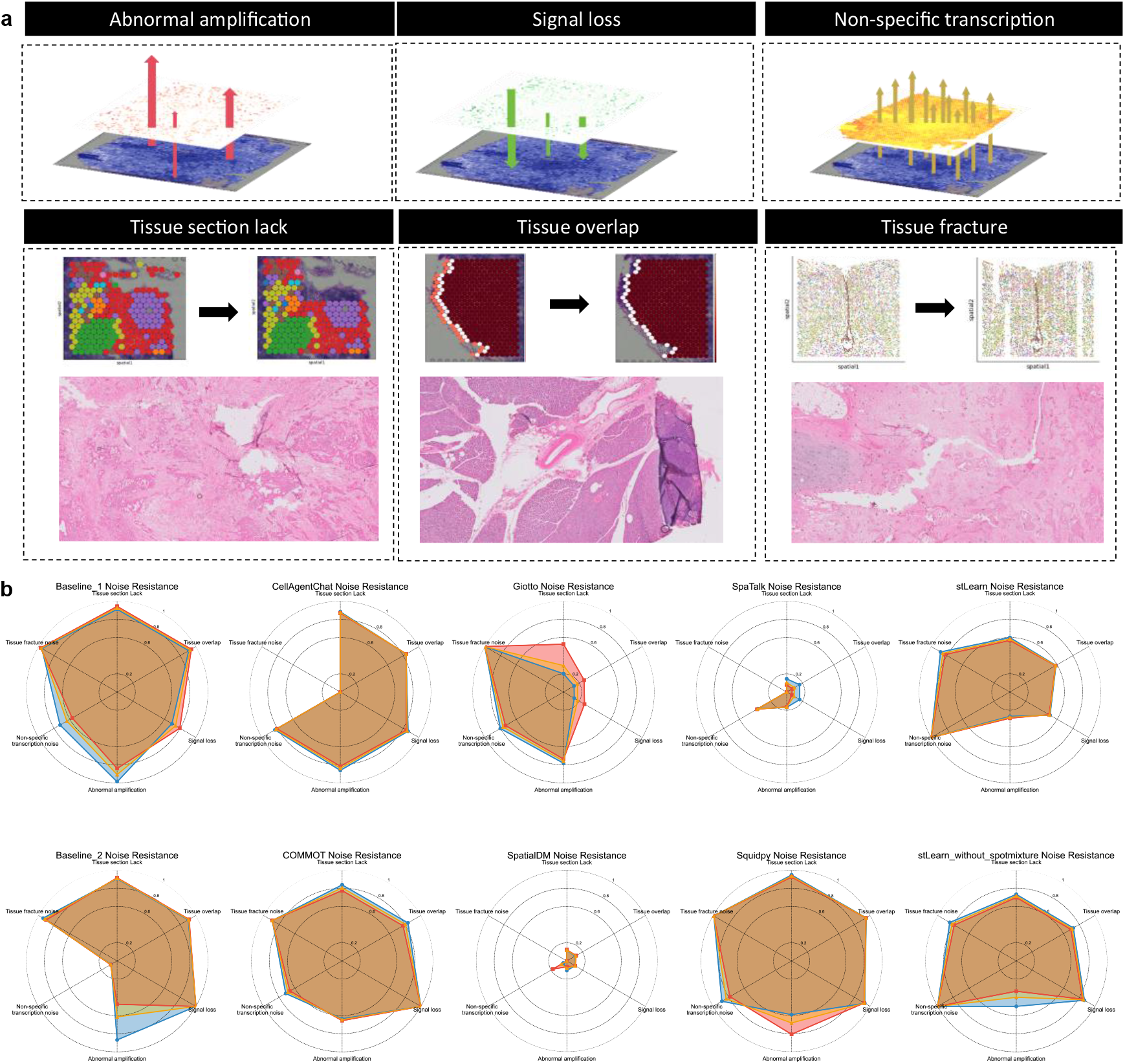
Robustness analysis under various noise conditions **a**. . Schematic diagram of noise simulation data construction. **b**. Evaluation of noise robustness of CCC tools under six simulated noises.

To assess anti-noise performance, the CCC results derived from the original data were treated as the ground truth, while those from noise-simulated data were used as observations to compute classification accuracy. As illustrated in Fig. 3b, Baseline 1 (based on neighbor permutation) exhibited slight CCC signal loss under spatial structure noise, whereas Baseline 2 (based on DEG analysis) was notably influenced by non-specific transcription noise. This indicates that both permutation- and DEG-dependent methods can be affected by technical and biological artifacts. However, their divergent failure modes highlight a methodological complementarity: permutation-based methods are more sensitive to spatial structure, whereas DEG-based methods are more sensitive to expression purity.

Among tools shown in Fig. 3b, COMMOT and Squidpy emerged as all-around performers, maintaining balanced radar profiles. In contrast, Giotto and CellAgentChat exhibited complementary robustness: Giotto excelled where coordinate precision was compromised, whereas CellAgentChat remained more stable during signal capture fluctuations. Tissue overlap and tissue fracture were identified as the most challenging scenarios, yielding the lowest average scores across all tools. This suggests that, although biochemical noise can be partially mitigated by specific algorithms, structural noise caused by technical errors remains a universal bottleneck for current spatial CCC inference. Specifically, stLearn showed the best resistance to non-specific transcriptional noise. Conversely, SpaTalk and SpatialDM were highly sensitive to noise, indicating their strong dependence on precise spot coordinates and accurate transcript detection.

### Overview of CCC analysis tools benchmarking and guidelines

To comprehensively compare the pros and cons of CCC analysis tools across the above analyses, we built an overall evaluation framework covering five dimensions: coverage, accuracy, spatial specificity, noise resistance, and user-friendliness.

As shown in Fig.4a, SpatialDM achieved the highest overall signal coverage, making it particularly suitable for rare CCC interaction identification, while COMMOT demonstrated superior classification accuracy and Squidpy stood out for its computational efficiency and effective pathway enrichment (see Fig. S4). Spatial specificity analysis revealed that stLearn is sensitive to spatial variance with high local-diversity, whereas SpatialDM’s dual Moran’s I module excels at identifying spatially autocorrelated signaling events. In terms of robustness, while Squidpy and COMMOT showed consistent stability across multiple noise types, SpatialDM and SpaTalk displayed higher sensitivity to biochemical noise such as signal loss.

**Fig. 4.**
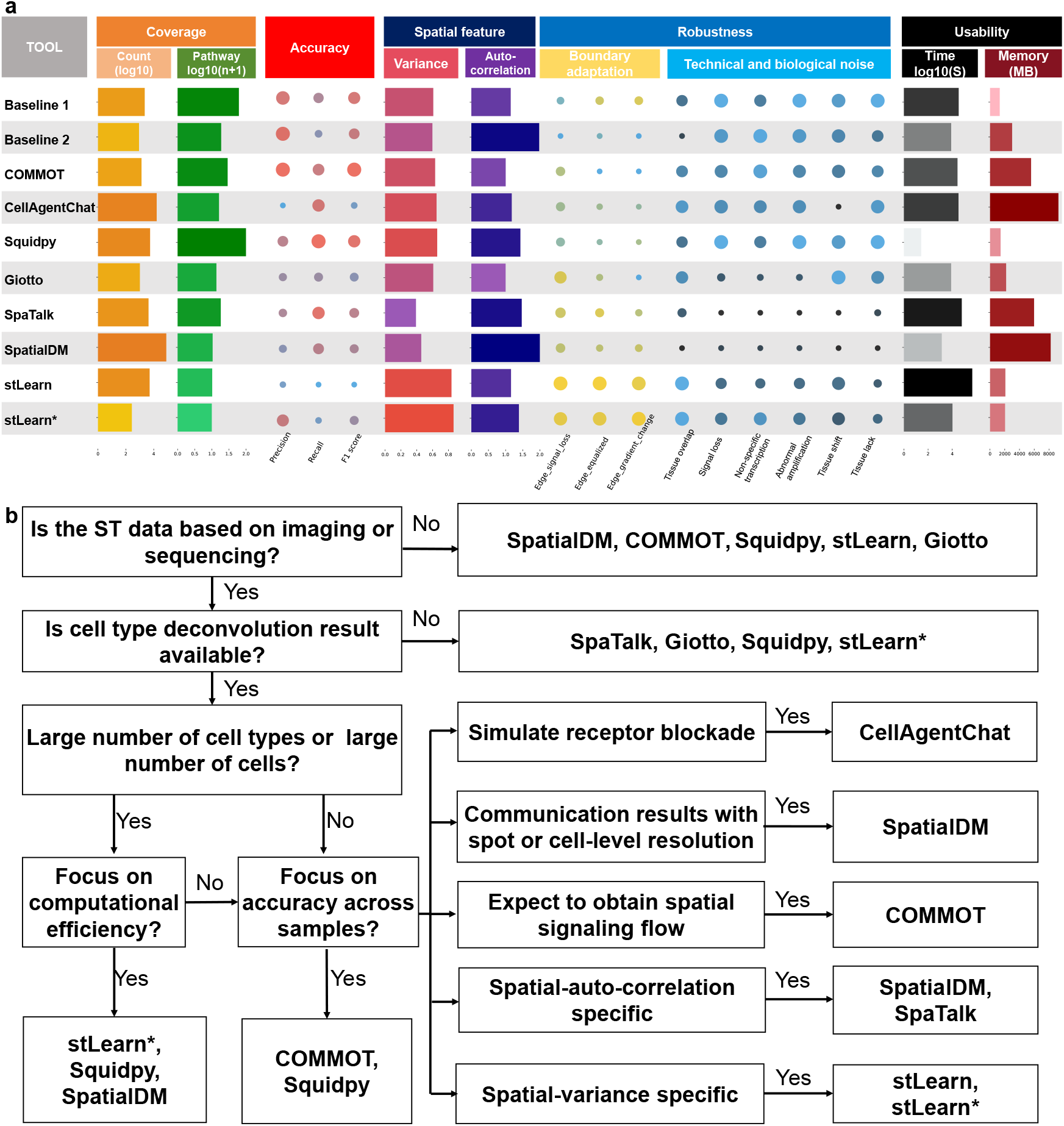
Overview of evaluation framework (a) and guidelines (b) for users (stLearn* indicates stLearn without spot-mixture mode). Coverage was assessed by LR-cell-cell counts and pathway gene enrichment. LR-cell-cell counts were averaged across 45 heterogeneous simulations, while pathway enrichment was based on ROI-associated gene counts (Detailed in Fig. S4). Accuracy was measured by precision, recall, and F1 score across samples (Detailed in Fig. S2b). Spatial variability was quantified by the mean normalized Simpson index across sampling ranges. Spatial autocorrelation was defined as the proportion of non-weak spatially autocorrelated interactions among all detected signals. Boundary applicability was evaluated using KS distance for edge equalization and reversed gradients, and mean signal loss across four edge-dropout conditions. Noise resistance was measured by F1 score before and after perturbation. Runtime and memory usage were averaged to assess user-friendliness (Detailed in Fig. S6).

Based on these performance profiles, we proposed a decision framework in Fig.4b to guide tool selection. This selection process begins with platform compatibility, where SpatialDM, COMMOT, Squidpy, stLearn, and Giotto are applicable to both imaging and sequencing data, and proceeds based on the availability of deconvolution results and the scale of the dataset, where Squidpy is recommended for its scalability. For specialized analytical goals, CellAgentChat is ideal for receptor blockade simulations, COMMOT for spatial signaling flow, and SpatialDM or stLearn for specific interests in autocorrelation or spatial variance, respectively. Ultimately, because no single tool excels across all biological contexts and ST data properties, we advocate for a goal-aligned, stepwise selection approach to ensure reliable and biologically interpretable CCC inference.

## Discussion

SpatialCCCbench provides a multi-dimensional benchmark for evaluating ST-based CCC inference tools across real datasets, simulated tissue partitions, noise perturbations and boundary-feature simulations. The results show that method performance depends strongly on platform, tissue structure, spatial resolution, cell-type annotation and noise type. Accuracy, coverage and robustness therefore need to be interpreted together rather than as isolated rankings.

Our assessment indicates that the heterogeneity and spatial specificity of ST data lead to complex CCC inference, with tissue structure and platform differences affecting accuracy. Among the tested methods, COMMOT, based on optimal transport distance, consistently achieved strong predictive performance, suggesting that spatial-distance-based models are effective for detecting widespread signaling. We also introduced a spatial autocorrelation framework in SpatialCCCbench, classifying four CCC signal types and evaluating spatial preference using Moran’s I and Geary’s C. Focusing on spatial MAUP[30], we quantified CCC signal diversity across spatial binning scales using Simpson’s index[29], showing that diversity generally decreases as sampling areas become smaller. However, some tools showed increased diversity at smaller scales, reflecting differences in sensitivity to resolution.

The continuous evolution of spatial technologies points toward the integration of multimodal data, such as spatial proteomics and metabolomics, to enhance CCC localization and molecular coverage[31-33]. Concurrently, advances in 3D tissue modeling and high-resolution capture technologies[34, 35] will enable the discovery of 3D cell-cell interactions within complex organoid architectures or intact tissue volumes, effectively capturing the volumetric constraints of cellular niches. As ST resolution approaches the single-cell or even subcellular level[36], mitigating stochastic gene loss and technical dropout becomes critical for maintaining the biological fidelity of interaction networks.

Moreover, integrating artificial intelligence with multi-omics data provides a transformative framework for constructing virtual cell models, where the precise characterization of CCC serves as a critical cornerstone[37-39]. Current virtual-cell studies are still largely characterized by CRISPR-based perturbation coupled with transcriptomic readouts[40], or by models trained only on scRNA-seq data[41]. Although these advances are valuable, they capture only one layer of cell state and therefore remain insufficient to comprehensively describe cellular mechanisms and behaviors. In reality, the coordinated actions of transcripts, proteins, and diverse metabolites more faithfully define more completely represent cellular states. A profound understanding of the tissue microenvironment, facilitated by spatially resolved multi-omics CCC analysis, will drive the development of virtual cell feature simulations in complex organisms, thereby supporting more reliable pathological interpretation and the advancement of precision medicine.

## Conclusions

SpatialCCCbench provides a multi-dimensional benchmark for evaluating ST-based CCC inference tools across real datasets, simulated tissue partitions, noise perturbations, and boundary-feature simulations. The benchmark demonstrates that method performance is strongly influenced by platform, tissue structure, spatial resolution, cell-type annotation, and noise type. Therefore, accuracy, coverage, spatial specificity, and robustness should be evaluated jointly rather than considered as independent rankings.

The results further indicate that the heterogeneity and spatial specificity of ST data create distinct analytical requirements for CCC inference. By integrating classification accuracy, signal coverage, biological relevance, spatial autocorrelation, local signal diversity, boundary-feature reliability, robustness, computational efficiency, and user-friendliness, SpatialCCCbench provides a standardized framework for characterizing the scenario-specific applicability of ST-based CCC tools. This framework supports systematic tool selection according to data modality, cell-type information, spatial resolution, noise characteristics, and analytical objectives.

Overall, SpatialCCCbench establishes a practical evaluation framework for ST-based CCC analysis and provides standardized metrics for interpreting CCC signals across diverse spatial biology contexts. This benchmark may facilitate the selection of appropriate CCC inference tools and support the development of future methods that better account for spatial heterogeneity, boundary features, and technical or biological noise.

## Methods

### Data preparation and preprocessing

We used multiple datasets in this study, including imaging-based and sequencing-based spatial transcriptomic data with or without spot-level cell-type mixture information. For classical spatial data with deconvoluted cell types, a Visium lymph node dataset paired with scRNA-seq data was deconvoluted using cell2location[42] to obtain spot-level cell-type composition. In addition, cluster labels were used as an alternative strategy for spot annotation in the DLPFC Visium and mouse brain MERFISH datasets, which do not explicitly preserve spot-mixture information. For these datasets, spot-mixture information was simulated by assigning the labeled cell type a fraction of 1 and all other cell types a fraction of 0. Raw data with cell-type annotations were quality controlled in Scanpy: genes were filtered using scanpy.pp.filter_genes (data, min_cells=1), total counts were normalized with scanpy.pp.normalize_total(data, target_sum=1e4), and mitochondrial, ribosomal, and hemoglobin genes were removed. The processed data were then organized as H5AD (Python format), st_count.csv (R format), and st_meta.csv (R format). LR reference information for CCC analysis was obtained from CellChatDB[43].

### Establishment of baseline

To systematically evaluate each tool’s ability to extract spatial information, we established two baselines to detect potential CCC signals in spatial transcriptomic data. Based on k-nearest neighbors (KNN), Baseline 1 identifies significant LR signals among the six nearest spots using the following permutation test:

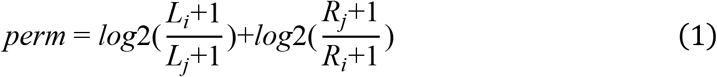

Here, L denotes ligand expression and R denotes receptor expression for sender spot i and receiver spot j. The values L_i_ and R_i_ are randomly shuffled to generate a background distribution, whereas L_j_ and R_j_ are kept fixed at their original values. A higher perm value indicates a stronger interaction between nearby spots, whereas perm values less than or equal to 0 indicate that ligand expression in the sender spot is lower than receptor expression in the same spot. Significant interactions in Baseline 1 were defined as high-ligand sender to high-receptor receiver pairs with permutation-derived P values < 0.05.

In addition to expression magnitude, Baseline 2 evaluates LR specificity through differential-expression analysis. A Student’s t-test was performed for receptor expression between the nearest spots (default: six neighbors around the sender) and distant spots outside the hypothesized diffusion area (default: spots at least 37 positions away), using sender spots with ligand expression > 0 as anchors. Ligands expressed in nearly all spots, for which distant reference spots could not be clearly defined, were treated as widely distributed genes without significant spatial interaction. Receptors with P values < 0.05 and log_2_FC > 0 were regarded as activated by the ligand in Baseline 2.

### Identifying cell-cell communication signals with different tools

Using these baselines, we were able to compare the signal preferences of different tools within the benchmarking framework. The implementation details for each tool are described below.

### Squidpy

Based on the Squidpy documentation (Documentation: https://squidpy.readthedocs.io/en/stable/api/squidpy.gr.ligrec.html#squidpy.gr.ligrec), the processed and quality-controlled H5AD file was read with scanpy.read_h5ad. squidpy.gr.ligrec was then used to detect communication among cell-type clusters with 1,000 permutations and CellChatDB as the interaction reference. LR pairs with P values < 0.05 were retained as significant interactions.

### CellAgentChat

Following the CellAgentChat tutorial in the project repository (Repository tutorial: https://github.com/mcgilldinglab/CellAgentChat/tree/main/tutorial), we built the CellAgentChat interaction model from processed spatial data and obtained interaction results with the CCI function using the retrained model. Because CellAgentChat requires integer row and column coordinates, it could not be directly applied to the MERFISH dataset, which lacks explicit row and column indices. Significant interactions were selected from p_value.csv using a threshold of P values < 0.05.

### COMMOT

Following the COMMOT documentation (Documentation: https://commot.readthedocs.io/en/latest/notebooks/visium-mouse_brain.html), genes in the LR reference were filtered using a minimum cell proportion of 0.05 in the spatial data. commot.tl.cluster_communication_spatial_permutation was then run with default parameters to infer CCC signals, and interactions with P values < 0.05 were retained.

### Giotto

Following the Giotto Suite documentation (Documentation: https://giottosuite.readthedocs.io/en/master/subsections/md_rst/specificCellCellcommunicationScores.html), a Giotto object was constructed from processed and quality-controlled CSV files containing cell and gene information. We used an additional function, spacomcell, to detect signals between each pair of cell types with default parameters. Significant interactions were defined using P values < 0.05, abs(log2FC) > 0.1, lig_nr >= 2, and rec_nr >= 2.

### SpaTalk

Following the SpaTalk tutorial (Documentation: https://raw.githack.com/multitalk/awesome-cell-cell-communication/main/method/tutorial.html), cell-type fractions and spot locations were reconstructed using generate_spot. CCC results were then obtained with dec_cci_all and filtered using the default threshold of P values < 0.05.

### SpatialDM

Following the SpatialDM documentation (Documentation: https://spatialdm.readthedocs.io/en/latest/melanoma.html), spatialdm.spatialdm_global was used to identify significant LR pairs through global Moran’s I, and spatialdm.spatialdm_local was used to identify significant local spot interactions. In addition, the ligand_ct and receptor_ct functions were used to obtain significant cell-type pairs. For consistency across methods, local cell-cell interactions with P values < 0.05 were defined as significant sender-receiver pairs.

### stLearn

Following the stLearn documentation (Documentation: https://stlearn.readthedocs.io/en/latest/tutorials/cell_cell_interaction.html), the LR reference was first processed with stlearn.tl.cci.run, and interaction results were obtained with stlearn.tl.cci.run_cci using the recommended parameters under both spot-mixture and no-spot-mixture modes. These two modes were treated as separate tool configurations in the benchmark because they use distinct input assumptions and produced substantially different outputs. Interactions with P values < 0.05 were retained.

### Spatial area splitting and noise simulation

Evaluating a tool’s performance requires more than a single dataset, and it necessitates benchmarks built on simulated data that faithfully represent the authentic complexities and requirements of spatial omics applications. To this end, we used the lymph node Visium data to perform area splitting and region of interest (ROI) extraction via SpatialData and napari[44, 45], followed by the simulation of six noise scenarios designed to test algorithmic resilience.

To systematically analyze spatial transcriptomics data at multiple resolutions, we implemented a partitioning algorithm that segments the tissue region of interest into progressively finer sub-rectangles. This approach facilitates the study of gene expression patterns across different spatial scales, enabling investigations into tool performance across multiple samples, applicability across different data scales, and the local signal diversity of cell-cell communication. To evaluate scale-dependent tool performance, we used SpatialData to generate multi-scale partitions from a rectangular tissue region of interest. The ROI was defined by its lower-left and upper-right coordinates and then subdivided into progressively finer spatial units across four scales, producing 45 subregions in total (3, 6, 12 and 24 subregions, respectively). Each subregion was exported as an independent H5AD file and subjected to the same quality-control, normalization and filtering workflow used for the original real-world datasets.

Specifically, six noise scenarios were simulated using real-world lymph node Visium data (see Fig. 3a). For transcript-level technical noise, abnormal amplification was modeled by randomly selecting 20% of genes in 20% of spots and doubling their expression values, whereas signal loss was modeled by randomly dropping out 20% of genes in 20% of spots. To model technical perturbations introduced during tissue preparation, we simulated tissue section lack and tissue overlap on two small tissue sections for computational efficiency. In these sections, selected spots were removed or edge-spot expression was merged separately. Tissue fracture noise was simulated on MERFISH data by moving the first 1,000 spots 100 pixels to the right and the last 1,000 spots 100 pixels to the left from their original positions. Background biological noise was modeled as non-specific transcriptional noise by adding Poisson-distributed noise corresponding to 10% of the existing gene expression counts.

### Benchmarking accuracy for classification of LR pairs maintaining CCC

To address the inherent heterogeneity in spatial transcriptomics data, where integrated signals from multiple samples may not reflect biologically meaningful ground truth due to inter-sample variability, this study focused on cell-cell communication signals mediated by ligand–receptor pairs within individual, non-integrated samples. For each sample, LR pairs identified by each tool were extracted. A consensus set of cell-cell interactions was established by considering only those interactions reported by at least three different tools as the ground truth. Baseline methods were included to ensure that the consensus captured nearest, specific, and significant interactions. The classification accuracy of each tool was then evaluated by calculating three standard metrics against this consensus ground truth as follows:

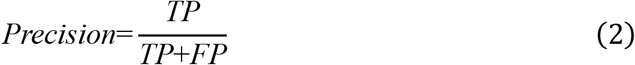

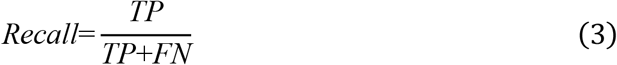

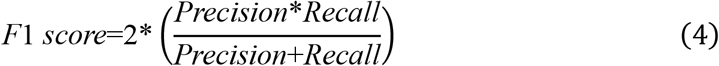

Finally, the mean values of three standard metrics across all evaluated samples were computed for each tool to provide an overall assessment of its predictive performance. As cell distribution and scale differ across samples, tools capture different signals due to detection preferences, and the accuracy of LR pairs fluctuates. To evaluate tool accuracy at the multi-sample level, ST data from split areas were processed to calculate standard metrics, and the mean values were used to assess the applicability of the tools.

### Benchmarking of pathway enrichment in ROI area

For ROI-focused pathway coverage analysis, follicle-enriched regions were extracted from the human lymph node Visium section using SpatialData, guided by the corresponding H&E image. The follicle ROI was extracted from the original lymph node dataset and saved as an independent H5AD object for identifying ROI-associated genes. Differential enrichment was assessed using Welch’s t-test for each gene, and genes with P < 0.05 and positive t statistics were retained as ROI-enriched genes. These genes were then subjected to KEGG pathway enrichment using Enrichr with the c2.cp.kegg_legacy.v2024.1.Hs.symbols.gmt gene-set collection. The significant KEGG pathways identified from the ROI-associated genes were used as reference pathways for pathway coverage analysis.

For each CCC inference method, ligand and receptor genes detected in the follicle ROI were extracted from the corresponding CCC results and subjected to the same KEGG enrichment workflow. Pathway coverage was quantified by calculating the gene ratio for each method across the ROI-associated KEGG pathways, with missing pathways assigned a value of zero. These pathway-level gene ratios were then summarized to compare the extent to which each method recovered ROI-associated signaling pathways.

### Benchmarking spatial autocorrelation of LR pairs

Considering that ligands and receptors exhibit characteristic distributions, four spatial autocorrelation LR interaction categories based on Moran’s I and Geary’s C were defined using two methods as follows.

For spatial autocorrelation analysis of individual genes in the interaction database, autocorrelation parameters from ESDA were designed as follows:

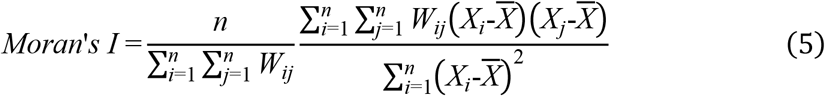

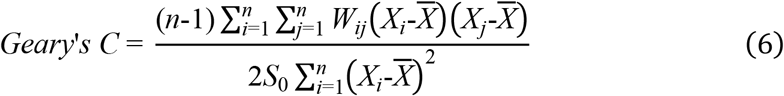

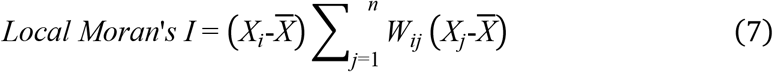

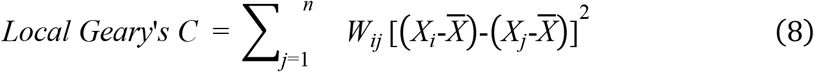

Here, X denotes the expression level of the ligand or receptor gene, W_ij_ denotes the spatial weight between spots i and j, and *S*_0_ is the sum of all spatial weights. For contact-dependent interactions, spatial weights were assigned to the six nearest neighbors, each with a value of 1. For diffusion-dependent interactions, spatial weights were assigned across 61 neighboring spots using a distance-decay scheme with six levels of weights: 10, 8, 6, 4, 2 and 0.

For LR-spatial correlation analysis, the expression levels of receptors and ligands were integrated into the same autocorrelation parameters, as follows:

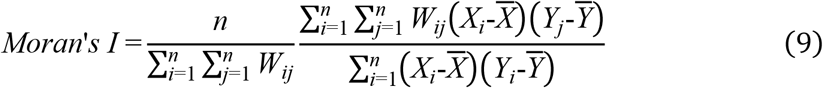

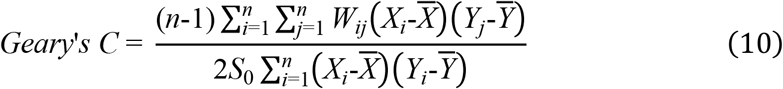

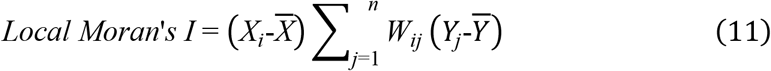

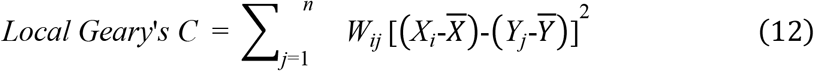

where X indicated the gene expression level of ligand, and Y indicated the gene expression of receptor. W implies the distance weight as defined above, and *S*_0_ indicate the sum of all weights.

Within the spatial autocorrelation framework, communication signals identified by each tool were classified into four categories: heterogeneous interaction LR, homogeneous interaction LR, random-like interaction LR, and weakly autocorrelated LR. First, widely distributed genes were filtered using the Gini index (default setting at 0.4) so that both ligand and receptor retained sufficient local spatial structure[46]. LR pairs with positive global Moran’s I values were then considered candidates for positive spatial autocorrelation, and local gene distributions were further evaluated using local Geary’s C and local Moran’s I, as described in Fig. S5a, b. Heterogeneous interaction LR pairs were defined by high local Geary’s C ≥ 1.5, and local Moran’s I > 0, representing expression differences around spots with high ligand expression and potential heterogeneous interaction. Homogeneous interaction LR pairs showed strong local aggregation, with local Moran’s I > 0 and local Geary’s C ∈(0, 0.5]. Random-like interaction LR pairs were defined by local Geary’s C ∈(0.5, 1.5) and global Moran’s I > 0. All remaining interactions were classified as weakly autocorrelated LR pairs.

### Benchmarking spatial variance of cell-cell communication signals in local areas

CCC signals can vary across spatial sampling scales. To quantify local signal diversity, we applied the Simpson index using the following formula:

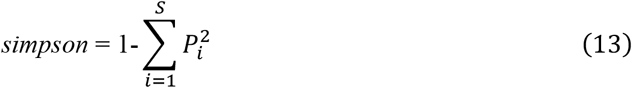

Here, P_i_ denotes the proportion of LR-cell-cell events assigned to the i-th LR pair among all detected LR-cell-cell events, and S denotes the total number of distinct LR pairs. The Simpson diversity index was calculated as 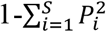 , where higher values indicate that interactions are more evenly distributed across LR pairs, whereas lower values indicate dominance by a small number of LR pairs.

### Benchmarking robustness of tools

To evaluate robustness, CCC signals inferred from the original data were compared with those inferred from simulated noisy data. LR-cell-cell pairs detected in the original data were treated as the ground truth, whereas pairs detected in the noisy data were treated as observations. As in the accuracy module, precision, recall, and F1 score were calculated, and overall anti-noise performance was summarized by the F1 score.

### Benchmarking boundary feature adaptation

To generate edges with distinct features for boundary feature adaptation analysis, we used a DLPFC dataset with clear cluster boundaries and specifically selected the interface between Layer 4 and Layer 5. As shown in Fig. S9, spots along this boundary were stratified into five spatial bands (Bands 1-5) and subjected to three types of artificial modification: edge equalization, signal loss through Poisson-based down sampling (intensities 0.2, 0.4, 0.6, and 0.8), and trans-gradient change.

The trans-gradient change was modeled using the constant-acceleration kinematic equation to simulate a nonlinear decay of gene expression across the tissue boundary.

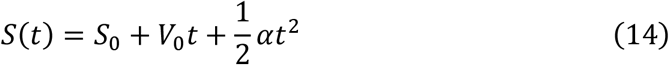

In this simulation, S(t) represents the cumulative reduction in expression relative to the initial state, S_0_ is the initial reduction at Band 1, V_0_ is the initial rate of reduction, and α is the constant deceleration term. In our simulation, S_0_ = 0%, V_0_ = 25%, and α = -6.25%, with t = 0, 1, 2, 3, and 4 corresponding to Bands 1-5. This yielded cumulative reductions of 0.000%, 21.875%, 37.500%, 46.875%, and 50.000%, corresponding to remaining expression levels of 100.000%, 78.125%, 62.500%, 53.125%, and 50.000%, respectively. This formulation provides a more biologically realistic simulation of boundary effects, in which ligand or receptor expression follows a graded transition rather than a simple linear decline.

Using these edge-feature simulations, we reran each tool on the perturbed datasets and compared the resulting CCC signals with those obtained from the original data, which served as the reference. The effect of different edge patterns was quantified by counting the number of receptors activated by each ligand. For edge equalization and trans-gradient simulations, the Pearson correlation between signal strengths in the original and perturbed datasets was used to assess edge adaptation, whereas the Kolmogorov-Smirnov (KS) distance between the original and noise-derived signal strength distributions was used to evaluate tool conservatism. For dropout intensities of 0.2, 0.4, 0.6, and 0.8, the 20 ligands with the highest receptor counts were selected to generate receptor-count curves and ligand rankings. Dependence on boundary gradients was then assessed by measuring curve fluctuation, calculated as the average deviation from the original result.

### Benchmarking computation efficiency on different scale data

Because spatial transcriptomic datasets are high-throughput and heterogeneous, sampling scale and cell-type complexity after deconvolution can strongly affect runtime and memory usage. When CCC analysis is performed across ROIs and deconvolution scales, both cell abundance and cell-type heterogeneity become important practical considerations. In addition, data dimensionality and sample size influence tool selection when computational time and hardware resources are limited. To evaluate computational efficiency across different scales, simulated datasets were divided into five groups ranging from the smallest ROI sample to the whole ST sample. Runtime for each tool was recorded using two central processing unit (CPU) cores and the default graphics processing unit (GPU) settings supported by each method, and peak memory usage was measured on the full real-world datasets with different cell type numbers.

Runtime and memory usage can also be affected by the number of cell types represented in the same spot set. For the lymph node Visium dataset, closely related subtypes from the same major lineage (for example, Macrophage_1 and Macrophage_2) were merged, generating four annotation scales with 7, 8, 10, and 17 major cell types, respectively, and the deconvolution fractions were updated accordingly. Runtime and memory usage were then recorded to assess tool performance under different levels of cell-type complexity.

All benchmarking analyses were performed on a Linux workstation running Ubuntu 18.04.6 LTS, equipped with an AMD Ryzen 5 5600 6-Core Processor (12 CPU threads available), and an NVIDIA GeForce RTX 3090 GPU with 24 GB of memory. Unless otherwise noted, runtime was benchmarked using two CPU cores together with the default GPU settings supported by each tool.

### Use of artificial intelligence (AI) tools

Large language models were used only for English-language editing and polishing of the manuscript text and for code organization and cleanup. No large language model was used for data analysis, result generation, figure production, or scientific interpretation. All authors reviewed and edited the final text and take full responsibility for the content of the manuscript.

## Declarations

## Ethics approval and consent to participate

This study used only publicly available, previously published, and fully anonymized datasets. No new human participants or human tissue were recruited or collected for this study, and no new animal experiments were performed. Ethics approval and informed consent were obtained by the original studies that generated the datasets, as described in the cited source publications. Therefore, no additional ethics approval was required for the analyses reported here.

## Availability of data and materials

The code used in this study is publicly available in the SpatialCCCbench repository (https://github.com/sgrazros/spatial-ccc-resource). The datasets used in this study are publicly available. The human DLPFC Visium dataset is available from Genome Research (DOI: https://doi.org/10.1101/gr.271288.120). The human lymph node Visium dataset and the matched reference single-cell dataset are available from the bioRxiv preprint by Kleshchevnikov et al. (DOI: https://doi.org/10.1101/2020.11.15.378125). The mouse brain MERFISH dataset was obtained from Moffitt et al. (https://www.science.org/doi/10.1126/science.aau5324). Additional simulated datasets are available in the SpatialCCCbench dataset repository (https://github.com/sgrazros/spatial-ccc-resource/Manuscript_Figure/datasets).

## Authors’ contributions

Q.W. and W.D. conceived and designed the research framework for the idea proposed by W.D.. Q.W. drafted the original manuscript and performed the benchmarking experiments. W.D. and D.L. contributed to manuscript review and proofreading. W.D. and Y.Y.L. supervised the study and secured funding. All authors read and approved the final manuscript.

## Acknowledgements

We appreciate the valuable feedback and suggestions from the early users of SpatialCCCbench. The authors acknowledge Beijing PARATERA Technology Co., LTD for providing high-performance and AI computing resources (http://cloud.paratera.com) that contributed to the research reported in this paper.

## References

1. Gulati GS, D'Silva JP, Liu Y, Wang L, Newman AM: Profiling cell identity and tissue architecture with single-cell and spatial transcriptomics. Nat Rev Mol Cell Biol 2025, 26:11–31.

2. Rao A, Barkley D, Franca GS, Yanai I: Exploring tissue architecture using spatial transcriptomics. Nature 2021, 596:211–220.

3. Zormpas E, Queen R, Comber A, Cockell SJ: Mapping the transcriptome: Realizing the full potential of spatial data analysis. Cell 2023, 186:5677–5689.

4. You Y, Fu Y, Li L, Zhang Z, Jia S, Lu S, Ren W, Liu Y, Xu Y, Liu X, et al: Systematic comparison of sequencing-based spatial transcriptomic methods. Nat Methods 2024, 21:1743–1754.

5. Jain S, Eadon MT: Spatial transcriptomics in health and disease. Nat Rev Nephrol 2024, 20:659– 671.

6. Oliveira MF, Romero JP, Chung M, Williams SR, Gottscho AD, Gupta A, Pilipauskas SE, Mohabbat S, Raman N, Sukovich DJ, et al: High-definition spatial transcriptomic profiling of immune cell populations in colorectal cancer. Nat Genet 2025, 57:1512–1523.

7. Jin Y, Zuo Y, Li G, Liu W, Pan Y, Fan T, Fu X, Yao X, Peng Y: Advances in spatial transcriptomics and its applications in cancer research. Mol Cancer 2024, 23:129.

8. Chelebian E, Avenel C, Wahlby C: Combining spatial transcriptomics with tissue morphology. Nat Commun 2025, 16:4452.

9. Sun C, Wang A, Zhou Y, Chen P, Wang X, Huang J, Gao J, Wang X, Shu L, Lu J, et al: Spatially resolved multi-omics highlights cell-specific metabolic remodeling and interactions in gastric cancer. Nat Commun 2023, 14:2692.

10. Hsieh WC, Budiarto BR, Wang YF, Lin CY, Gwo MC, So DK, Tzeng YS, Chen SY: Spatial multi-omics analyses of the tumor immune microenvironment. J Biomed Sci 2022, 29:96.

11. Wu Y, Cheng Y, Wang X, Fan J, Gao Q: Spatial omics: Navigating to the golden era of cancer research. Clin Transl Med 2022, 12:e696.

12. Cesaro G, Nagai JS, Gnoato N, Chiodi A, Tussardi G, Kloker V, Musumarra CV, Mosca E, Costa IG, Di Camillo B, et al: Advances and challenges in cell-cell communication inference: a comprehensive review of tools, resources, and future directions. Brief Bioinform 2025, 26:bbaf280.

13. Walker BL, Cang Z, Ren H, Bourgain–Chang E, Nie Q: Deciphering tissue structure and function using spatial transcriptomics. Commun Biol 2022, 5:220.

14. Peng L, Wang F, Wang Z, Tan J, Huang L, Tian X, Liu G, Zhou L: Cell-cell communication inference and analysis in the tumour microenvironments from single-cell transcriptomics: data resources and computational strategies. Brief Bioinform 2022, 23:bbac234.

15. Pham D, Tan X, Balderson B, Xu J, Grice LF, Yoon S, Willis EF, Tran M, Lam PY, Raghubar A: Robust mapping of spatiotemporal trajectories and cell–cell interactions in healthy and diseased tissues. Nature communications 2023, 14:7739.

16. Li Z, Wang T, Liu P, Huang Y: SpatialDM for rapid identification of spatially co-expressed ligand-receptor and revealing cell-cell communication patterns. Nat Commun 2023, 14:3995.

17. Zhu J, Wang Y, Chang WY, Malewska A, Napolitano F, Gahan JC, Unni N, Zhao M, Yuan R, Wu F, et al: Mapping cellular interactions from spatially resolved transcriptomics data. Nat Methods 2024, 21:1830–1842.

18. Cang Z, Zhao Y, Almet AA, Stabell A, Ramos R, Plikus MV, Atwood SX, Nie Q: Screening cell-cell communication in spatial transcriptomics via collective optimal transport. Nat Methods 2023, 20:218–228.

19. Yuan Y, Bar–Joseph Z: GCNG: graph convolutional networks for inferring gene interaction from spatial transcriptomics data. Genome Biol 2020, 21:300.

20. Raghavan V, Zheng Y, Li Y, Ding J: Harnessing agent-based frameworks in CellAgentChat to unravel cell–cell interactions from single-cell and spatial transcriptomics. Genome Research 2025, 35:1646–1663.

21. Armingol E, Baghdassarian HM, Lewis NE: The diversification of methods for studying cell-cell interactions and communication. Nat Rev Genet 2024, 25:381–400.

22. Plummer JT, Dezem FS, Cook DP, Park J, Zhang L, Liu Y, Marcao M, DuBose H, Wani A, Wise K, et al: Standardized metrics for assessment and reproducibility of imaging-based spatial transcriptomics datasets. Nat Biotechnol 2025. doi:10.1038/s41587-025-02811-9.

23. You Y, Fu Y, Li L, Zhang Z, Jia S, Lu S, Ren W, Liu Y, Xu Y, Liu X, et al: Systematic comparison of sequencing-based spatial transcriptomic methods. Nature Methods 2024, 21:1743–1754.

24. Liu Z, Sun D, Wang C: Evaluation of cell-cell interaction methods by integrating single-cell RNA sequencing data with spatial information. Genome Biol 2022, 23:218.

25. Luo J, Deng M, Zhang X, Sun X: ESICCC as a systematic computational framework for evaluation, selection, and integration of cell-cell communication inference methods. Genome Res 2023, 33:1788–1805.

26. Chen JG, Chávez–Fuentes JC, O’Brien M, Xu J, Ruiz EC, Wang W, Amin I, Sheridan JP, Shin SC, Hasyagar SV, et al: Giotto Suite: a multiscale and technology-agnostic spatial multiomics analysis ecosystem. Nature Methods 2025, 22:2052–2064.

27. Anselin L: Local Indicators of Spatial Association—LISA. Geographical Analysis 1995, 27:93– 115.

28. Rey SJ, Anselin L: PySAL: A Python library of spatial analytical methods. In Handbook of applied spatial analysis: Software tools, methods and applications. Springer; 2009: 175–193

29. Nagendra H: Opposite trends in response for the Shannon and Simpson indices of landscape diversity. Applied Geography 2002, 22:175–186.

30. Ding DY, Tang Z, Zhu B, Ren H, Shalek AK, Tibshirani R, Nolan GP: Quantitative characterization of tissue states using multiomics and ecological spatial analysis. Nature Genetics 2025, 57:910–921.

31. Liu L, Li W, Wang F, Li Y, Huang LK, Wong KC, Yang F, Yao J: A pre-trained large generative model for translating single-cell transcriptomes to proteomes. Nat Biomed Eng 2025. doi:10.1038/s41551-025-01528-z.

32. Wagner M, Kang J, Mercado C, Thirumlaikumar VP, Gorka M, Zillmer H, Guo J, Minen RI, Plecki CF, Dehesh K, et al: Mapping protein-metabolite interactions in E. coli by integrating chromatographic techniques and co-fractionation mass spectrometry. iScience 2025, 28:112611.

33. Lee KS, Su X, Huan T: Metabolites are not genes -avoiding the misuse of pathway analysis in metabolomics. Nat Metab 2025, 7:858–861.

34. Qiu X, Zhu DY, Yao J, Jing Z, Zuo L, Wang M, Min KH, Pan H, Wang S, Liao S, et al: Spateo: multidimensional spatiotemporal modeling of single-cell spatial transcriptomics. bioRxiv 2022:2022.2012.2007.519417.

35. Kern C, Zhang Q, Lu Y, Eschbach J, Zeng Z, Farah EN, Tai C-Y, Yang K, Jenie I, Yao F, et al: MERFISH+, a large-scale, multi-omics spatial technology resolves the molecular holograms of the 3D human developing heart. bioRxiv 2025:2025.2011.2002.686137.

36. Zhao Y, Li Y, He Y, Wu J, Liu Y, Li X, Li Z, Yuan Q, Li J, Zhang X, et al: Stereo-seq V2: Spatial mapping of total RNA on FFPE sections with high resolution. Cell 2025, 188:6554–6571 e6521.

37. Cui H, Wang C, Maan H, Pang K, Luo F, Duan N, Wang B: scGPT: toward building a foundation model for single-cell multi-omics using generative AI. Nat Methods 2024, 21:1470–1480.

38. Zhang H, Yuan G-H, Yuan C, Xu T, Bian T, Cheng H, Huang W, Zhao D, Rong Y: Lingshu-Cell: A generative cellular world model for transcriptome modeling toward virtual cells. arXiv preprint arXiv:260325240 2026.

39. Noutahi E, Hartford J, Tossou P, Whitfield S, Denton AK, Wognum C, Ulicna K, Craig M, Hsu J, Cuccarese M: Virtual cells: Predict, explain, discover. arXiv preprint arXiv:250514613 2025.

40. Wang C, Karimzadeh M, Ravindra NG, Bounds LR, Alerasool N, Huang AC, Ma S, Gulbranson DR, Cui H, Lee Y, et al: X-Cell: Scaling Causal Perturbation Prediction Across Diverse Cellular Contexts via Diffusion Language Models. bioRxiv 2026:2026.2003.2018.712807.

41. Gandhi S, Javadi F, Svensson V, Khan U, Jones MG, Yu J, Merico D, Goodarzi H, Alidoust N: Tahoe-x1: Scaling Perturbation-Trained Single-Cell Foundation Models to 3 Billion Parameters. bioRxiv 2025:2025.2010.2023.683759.

42. Kleshchevnikov V, Shmatko A, Dann E, Aivazidis A, King HW, Li T, Elmentaite R, Lomakin A, Kedlian V, Gayoso A, et al: Cell2location maps fine-grained cell types in spatial transcriptomics. Nat Biotechnol 2022, 40:661–671.

43. Jin S, Plikus MV, Nie Q: CellChat for systematic analysis of cell-cell communication from single-cell transcriptomics. Nat Protoc 2025, 20:180–219.

44. Marconato L, Palla G, Yamauchi KA, Virshup I, Heidari E, Treis T, Vierdag W-M, Toth M, Stockhaus S, Shrestha RB, et al: SpatialData: an open and universal data framework for spatial omics. Nature Methods 2025, 22:58–62.

45. Ahlers J, Althviz Moré D, Amsalem O, Anderson A, Bokota G, Boone P, Bragantini J, Buckley G, Burt A, Bussonnier M: napari: a multi-dimensional image viewer for Python. Zenodo 2023. doi:10.5281/zenodo.8115575.

46. Friel S, Akerman M, Hancock T, Kumaresan J, Marmot M, Melin T, Vlahov D: Addressing the social and environmental determinants of urban health equity: evidence for action and a research agenda. J Urban Health 2011, 88:860–874.

