## Supplemental Figure S1-S9 for "SpatialCCCbench: Standardized Metrics for the Systematic Evaluation of Spatial Cell-Cell Communication Methods"

### Supplementary Materials

This supplementary material provides additional figures and analyses supporting the main manuscript, including benchmark performance across datasets, coverage and similarity of ligand-receptor interactions, enrichment analysis, spatial autocorrelation-based interaction classification, user-friendliness evaluation, and summary comparisons across tools.

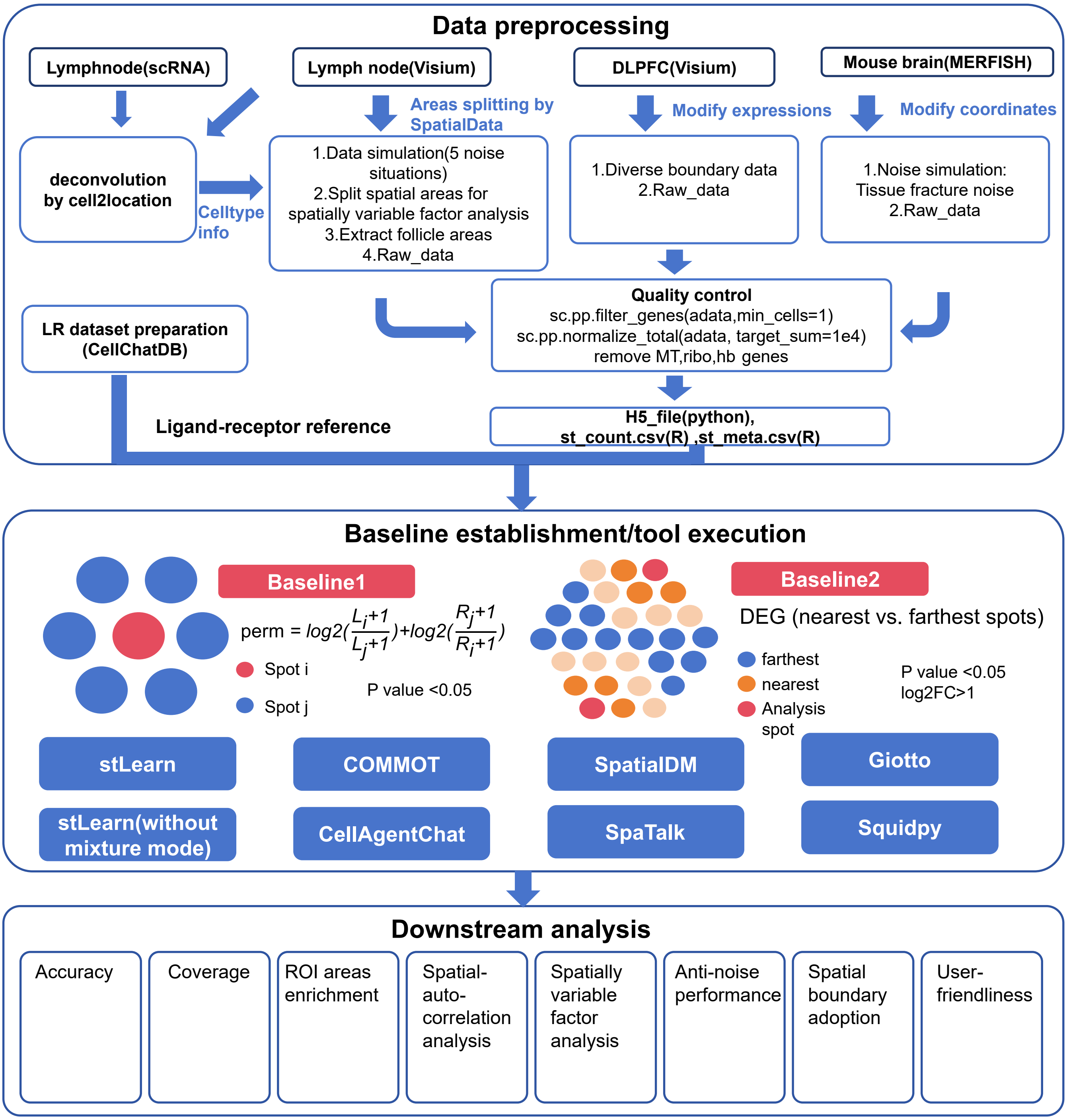


Figure S1 Workflow of SpatialCCCbench

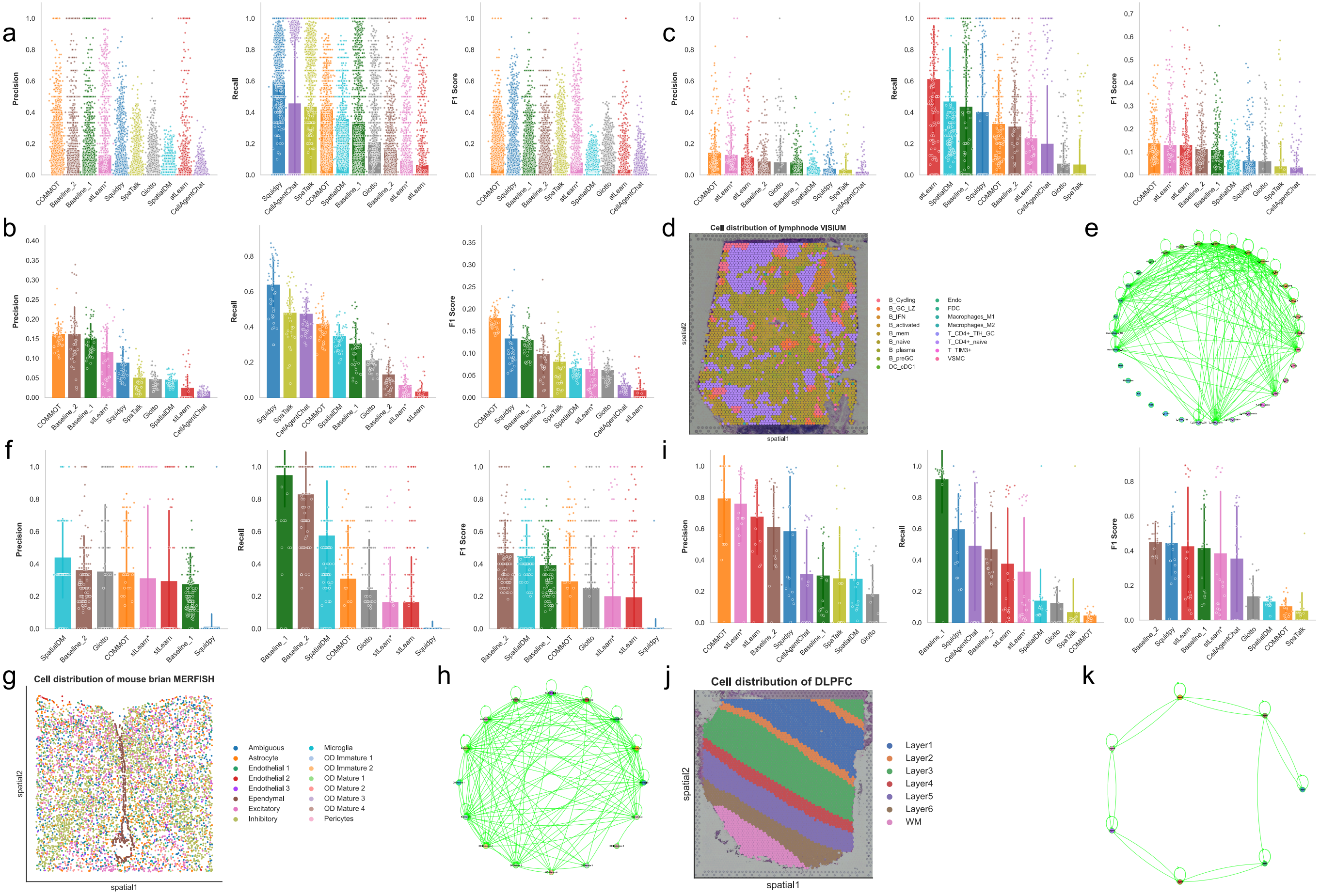


Figure S2 Classification accuracy of CCC signals across all samples and individual samples

a. Accuracy of LR-pair predictions across different cell-cell interactions in all samples. Sample-level accuracy is not evaluated in this panel; each point represents the prediction accuracy of LR-pairs within one cell-cell interaction.

b. Accuracy evaluation of LR-cell-cell interactions across individual samples. Each point represents one sample source and reflects sample-level evaluation. c-k. Evaluation across three real-world datasets: human lymph node (c-e), mouse brain MERFISH (f-h), and human DLPFC (i-k), including classification accuracy (c, f, i), cell-type distributions (d, g, j), and the ground-truth interactions used for validation (e, h, k). SpaTalk and CellAgentChat were excluded from the accuracy evaluation on the MERFISH dataset because SpaTalk requires spot reconstruction and CellAgentChat requires integer-valued spot coordinates. In the network plots in e, h, and k, nodes represent cell types, edges represent interacting ligand-receptor (LR) pairs, and edge thickness indicates the number of interacting LR pairs.

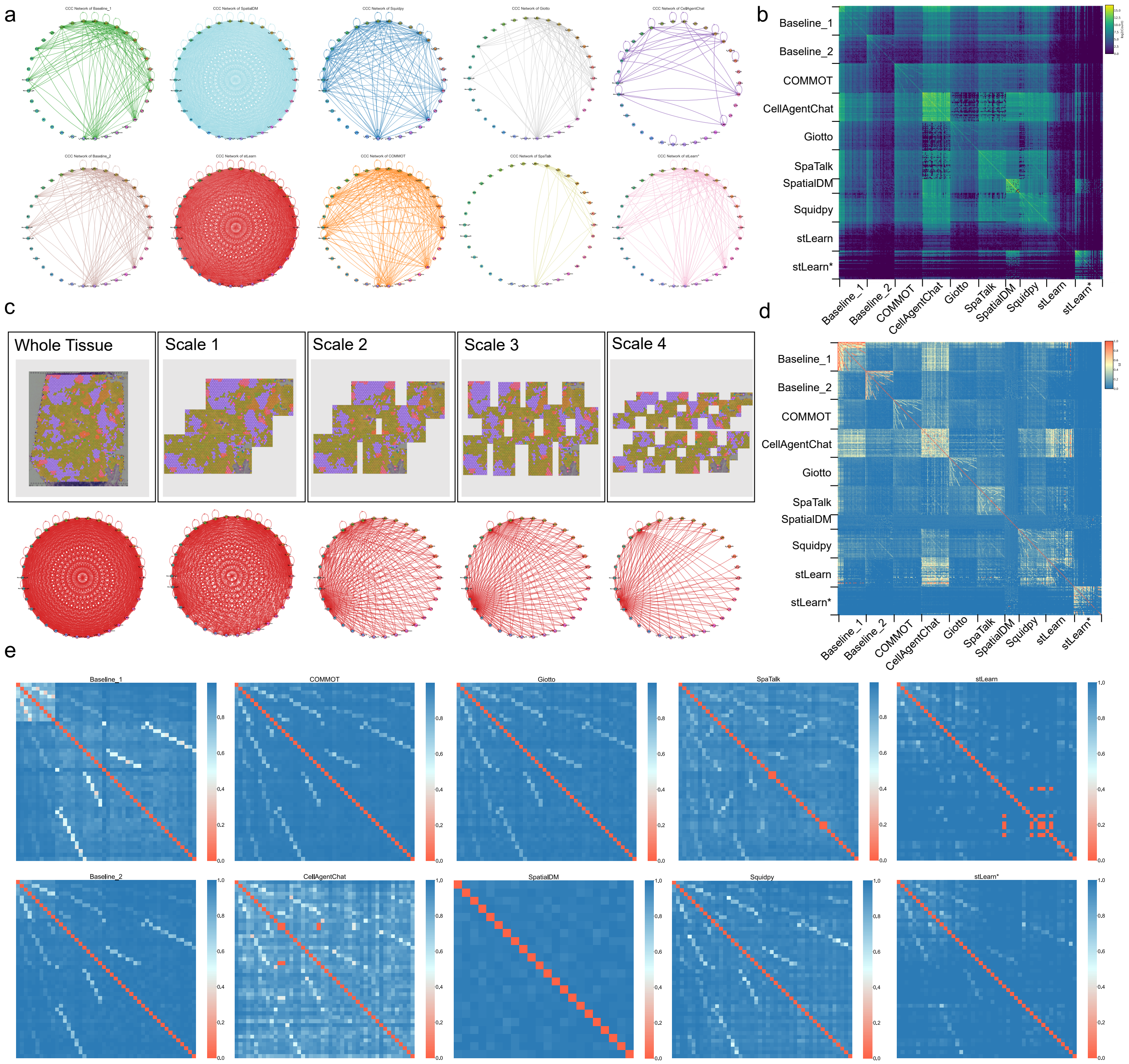


Figure S3 Coverage and similarity of LR pairs associated with CCC across tools

a. CCC interaction networks reconstructed by different tools in the 10x Visium lymph node dataset.

b, d. Heatmaps depicting the counts of intersecting LR-cell-cell pairs (b) and the similarity index (d) across multiple samples for the evaluated CCC tools.

c. Trends in CCC signals across different sample scales.

e. Heatmaps depicting the Jaccard distance across multiple samples for each individual tool.

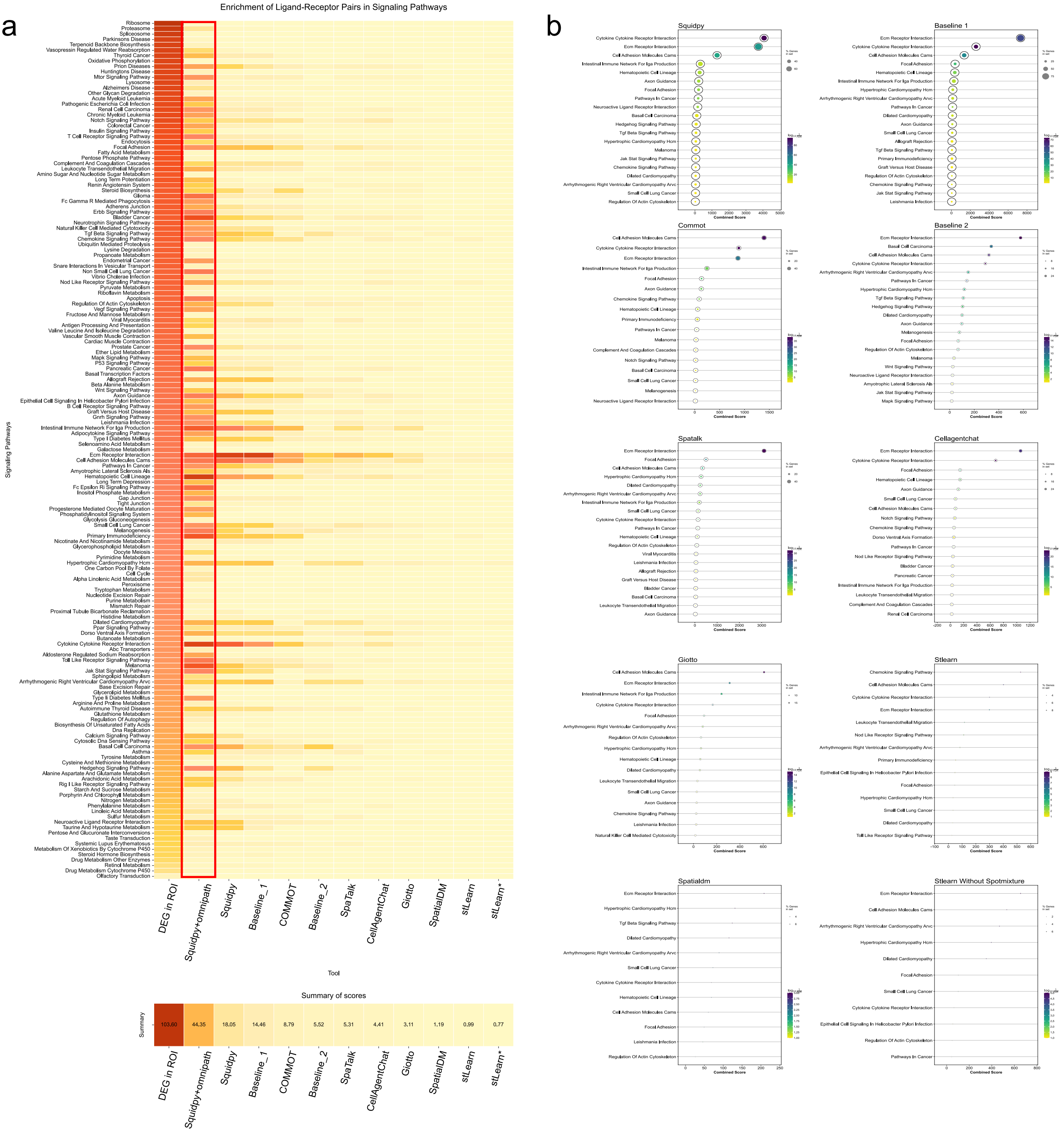


Figure S4 Enrichment analysis reveals distinct patterns and network attributes of DEGs and LR genes across tools.

a. Patterns of pathway enrichment of DEGs from follicles and LR genes identified by the evaluated CCC tools.

b. Pathway enrichment and statistical attributes of CCC signals for each tool.

#### Enrichment of signals and biological effects in ROIs for CCC tools

ST enables localized tissue exploration via region-of-interest (ROI) analysis. In the signal enrichment analysis of Visium lymph node follicles shown in Fig. S4a, DEG analysis identified protein synthesis pathways, while CCC tools captured diverse communication-related pathways such as ECM-receptor interactions and cell adhesion.

Tool-specific preferences were evident in Fig. S4a, b. Squidpy, integrated with the continuously updated OmniPath database, achieved the most comprehensive ROI-specific enrichment of signaling patterns. It uniquely identified T/B cell receptor signaling aligned with immune function, likely due to its permutation-based testing and database breadth. Conversely, distance-constrained methods focused on focal adhesion, while stLearn excelled at identifying soluble signals such as chemokines.

Despite these insights, many CCC signals lacked prominent KEGG enrichment. This highlights a gap in downstream pathway annotation, necessitating further exploration to fully decode complex intercellular networks within specific spatial ROIs.

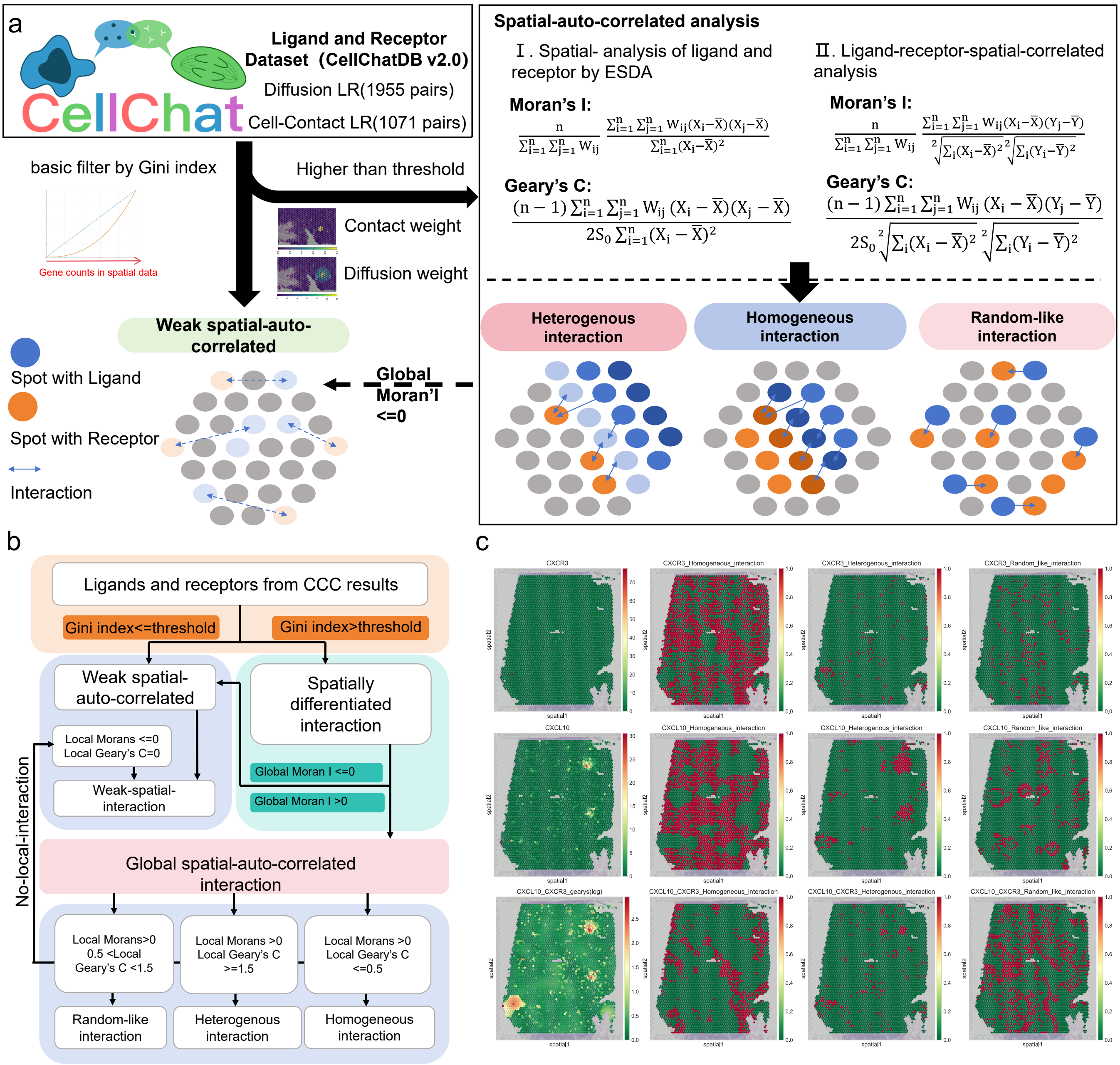


Figure S5 Framework for analyzing ligand-receptor interactions based on spatial autocorrelation.

a. Framework for analyzing ligand-receptor interactions based on spatial autocorrelation.

b. Workflow for interaction classification based on spatial autocorrelation analysis.

c. Effects of four spatial autocorrelation features in real spatial transcriptomics data.

#### Interpretation of spatial autocorrelation patterns

The interaction classification scheme is based on Global Moran's I, Local Moran's I, and Local Geary's C. For spatial autocorrelation features of individual genes, either the ligand or the receptor must exhibit the corresponding spatial autocorrelation feature. For LR-integrated spatial autocorrelation features, the autocorrelation parameters calculated from LR integration must satisfy the relevant classification criteria. Using CXCL10-CXCR3 as an example and referring to the original gene distribution, the locally heterogeneous autocorrelation pattern can characterize signal diffusion changes within the interaction region; when the Local Geary's C threshold is set to 1.5, it better captures ligand diffusion. In regions with locally homogeneous autocorrelation, ligand and receptor distributions are relatively even. Regions with random-like local autocorrelation, where Local Geary's C is close to 1, exhibit a random distribution that is neither clustered nor uniform.

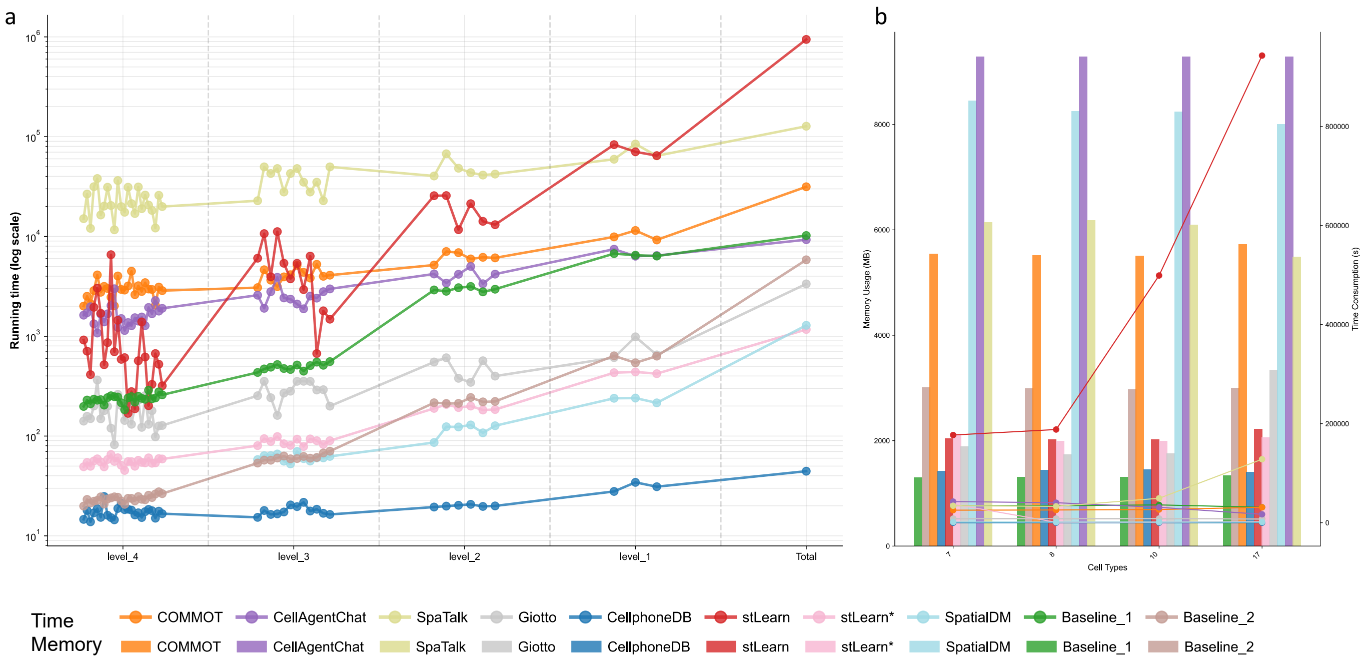


Figure S6 User-friendliness analysis of tools across different cell numbers and cell types

a. Running time comparisons across tools for different cell numbers at different sampling scales.

b. Running time and memory usage comparisons across tools for different cell types at different sampling scales.

#### User-friendliness analysis of tools across different cell numbers and cell types

The rapid evolution of GPU hardware capabilities, coupled with continuous innovations in high-performance computing frameworks, has significantly enhanced the efficiency of deep learning model training and inference. However, fully harnessing this hardware potential critically depends on the optimization and usability of software toolchains. Tools that maintain low memory and CPU overhead exhibit broader applicability across diverse hardware environments. Not all research laboratories possess high-performance GPU clusters. Therefore, tools with optimized, lightweight architectures can be seamlessly deployed on standard workstations or personal laptops.

In this study, the user-friendliness of the tools was evaluated by recording runtime and memory usage across 46 spatial datasets with varying numbers of cell types and spots, categorized into five scales as illustrated in Fig. S6a. Under consistent hardware conditions detailed in the Methods, both runtime and memory consumption were found to be strongly correlated with the number of spots and cell types.

As shown for runtime in Fig. S6a, all tools completed analyses within 10^5 seconds on small-scale datasets, while Squidpy demonstrated superior computational efficiency and SpaTalk required the longest processing time. In the original large-scale spatial transcriptomic data containing thousands of spots, stLearn and SpaTalk exhibited substantially increased runtimes, whereas Squidpy maintained efficient utilization of computing resources, thereby demonstrating high scalability. Given the high density and heterogeneity of spatial transcriptomic data, a trade-off between computational efficiency and analytical accuracy appears inevitable.

In terms of memory usage (Fig. S6b), tools that maintain low memory and CPU overhead, referring here to the peak total memory generated across both CPU and GPU during execution, exhibit broader applicability across diverse hardware environments. As shown in Fig. S6b, CellAgentChat required the highest memory allocation, with peak usage across CPU and GPU exceeding 8 GB. This intensive consumption can be attributed to its complex process of model reconstruction and the integration of high-dimensional feature vectors derived from spatial data. In contrast, Squidpy again exhibited low memory requirements, highlighting its high scalability.

Furthermore, the number of cell types also influenced computational resource consumption, with variation among tools due to differences in methodological principles. According to Fig. S6b, memory usage was only mildly affected by the number of cell types. In contrast, datasets with more cell types consistently led to longer runtimes, suggesting that cluster size and cellular diversity significantly affected the computational performance of the tools. Notably, tools such as SpaTalk, CellAgentChat, and stLearn represent the high-demand extreme, whereas Squidpy serves as a highly accessible, low-resource alternative, highlighting the importance of selecting tools based on the specific balance of available hardware and required analytical resolution.

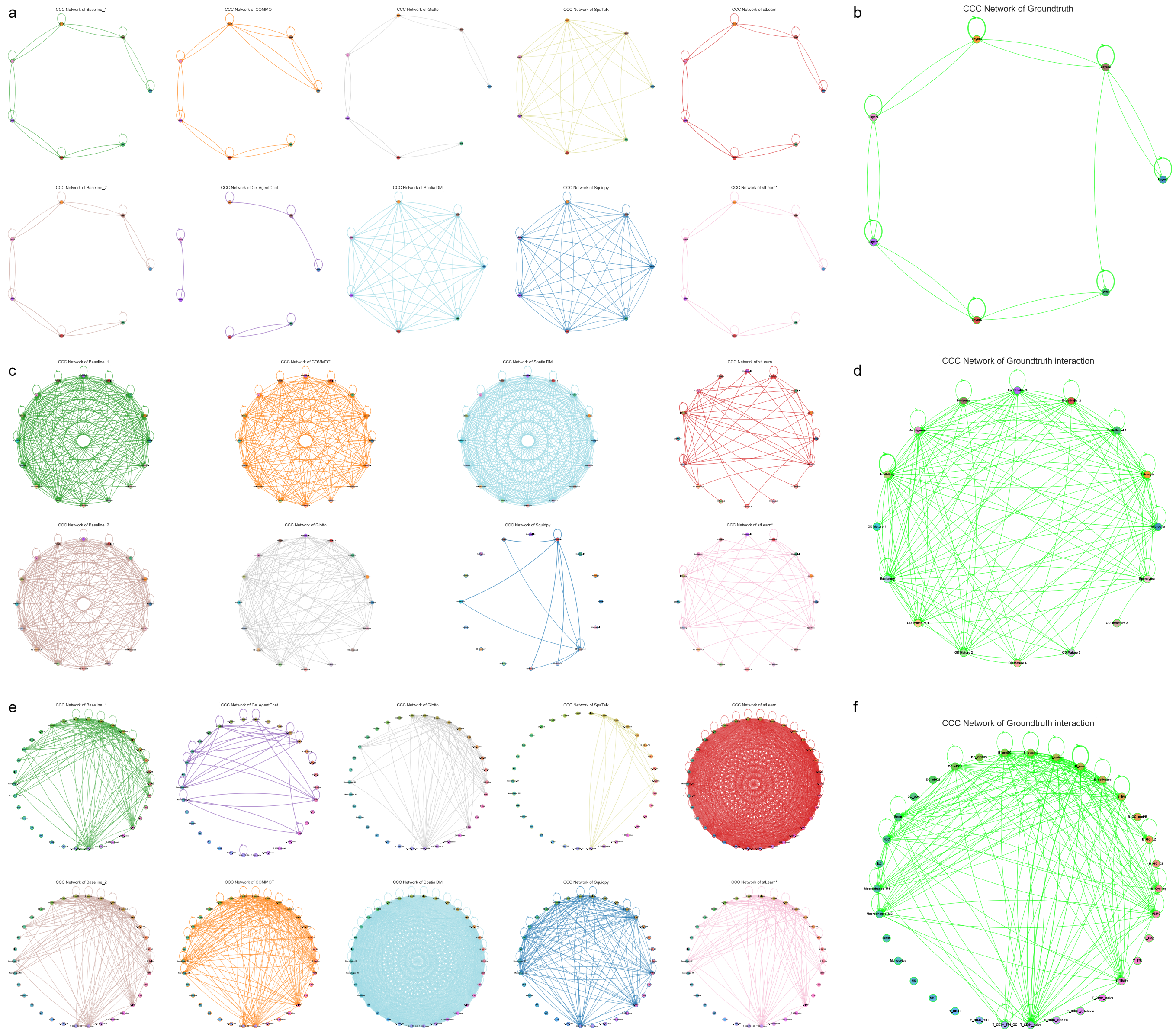

Figure S7 Summary of CCC networks from different datasets.

a, c, e. CCC interaction networks reconstructed by different tools for DLPFC Visium, mouse brain MERFISH, and lymph node Visium samples.

b, d, f. Ground-truth CCC signals for DLPFC Visium, mouse brain MERFISH, and lymph node Visium samples.

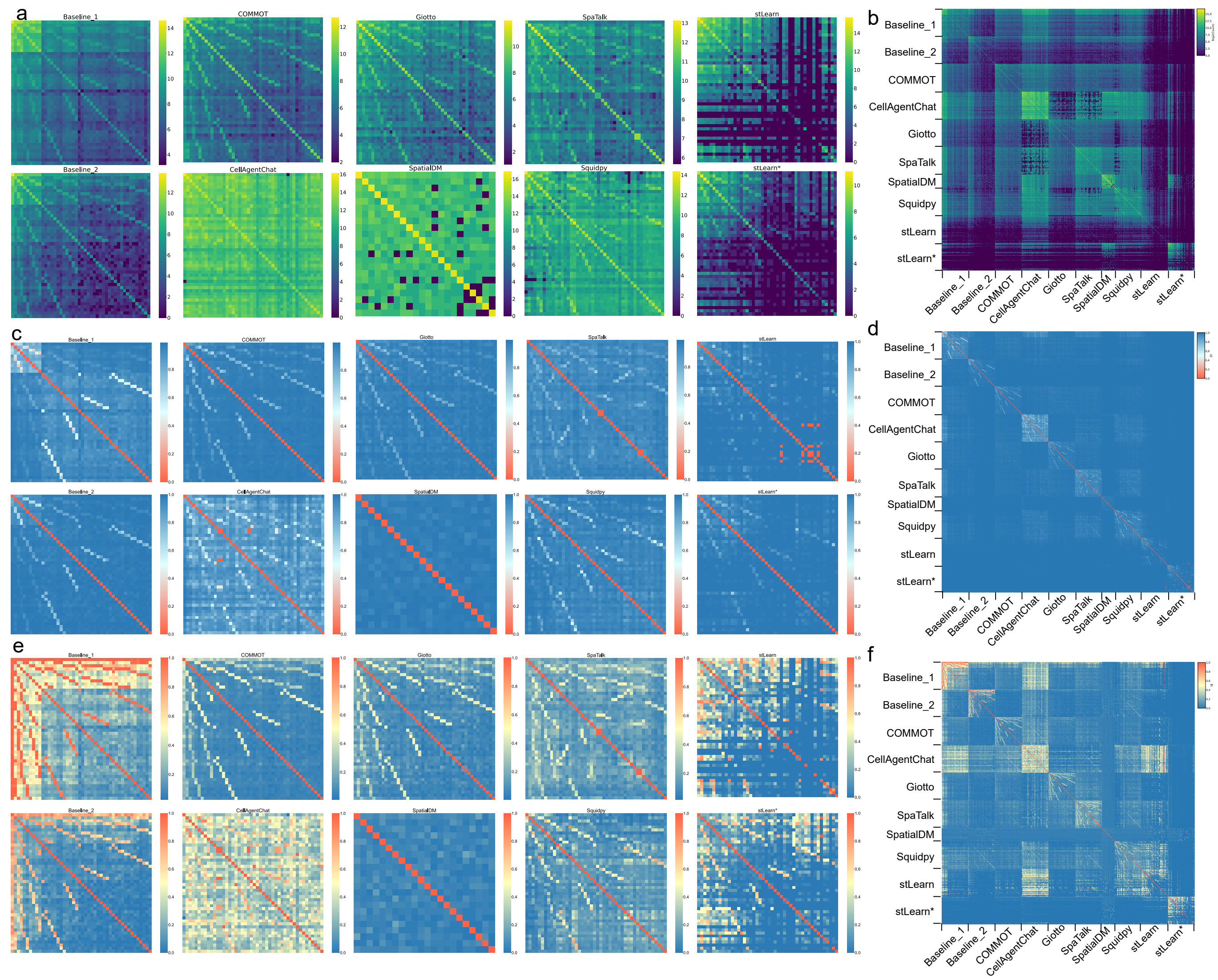


Figure S8 Summary of similarity analyses across tools.

a, c, e. Heatmaps depicting the counts of intersecting LR-cell-cell pairs (a), Jaccard distance (c), and similarity index (e) across multiple samples for each individual tool.

b, d, f. Heatmaps depicting the counts of intersecting LR-cell-cell pairs (b), Jaccard distance (d), and similarity index (f) across multiple samples for the evaluated CCC tools. For each tool in Figure S8, the multiple samples are ordered from highest to lowest spot number.

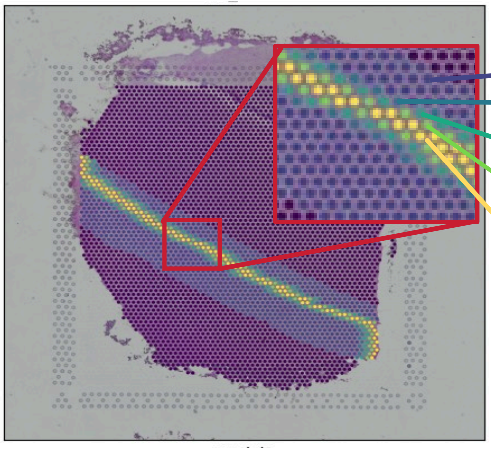

| Gradient change | Edge equalization | Edge signal dropout noise |
|---|---|---|
| Original expression | Average total gene expression across bands 1–5 | Poisson-based down-sampling of cells from regions 1–5 (intensities: 0.2, 0.4, 0.6, 0.8). |
| Expressed at 78.125% of the original | | |
| Expressed at 62.500% of the original | | |
| Expressed at 53.125% of the original | | |
| Expressed at 50% of the original | | |

Gradient change is designed by

$$S(t) = S_0 + V_0 t + \frac{1}{2}\alpha t^2$$

Figure S9 Simulation design of spatial signal gradients.
